# An Atlas of Protein Phosphorylation Dynamics During Interferon Signaling

**DOI:** 10.1101/2024.06.28.601147

**Authors:** Idoia Busnadiego, Marie Lork, Sonja Fernbach, Samira Schiefer, Nikos Tsolakos, Benjamin G. Hale

## Abstract

Interferons (IFNs, types I-III) have pleiotropic functions in promoting antiviral and anti-tumor responses, as well as in modulating inflammation. Dissecting the signaling mechanisms elicited by different IFNs is therefore critical to understand their phenotypes. Here, we use mass spectrometry to investigate the early temporal dynamics of cellular protein phosphorylation in response to stimulation with IFNα2, IFNβ, IFNω, IFNγ, and IFNλ1, representing all IFN types. We report an atlas of over 700 common or unique phosphorylation events reprogrammed by these IFNs, revealing both previously known and uncharacterized modifications. Functional screening and mechanistic studies identify that several factors differentially-modified in response to IFNs contribute to host antiviral responses, either directly or by supporting IFN-stimulated gene or protein production. Among these, phosphorylation of PLEKHG3 at serine-1081 creates a phospho-regulated binding motif for the docking of 14-3-3 proteins, and together these factors contribute to coordinating efficient IFN-stimulated gene expression independent of early JAK/STAT signaling. Our findings map the global phosphorylation landscapes regulated by IFN types I, II, and III, and provide a key resource to explore their functional consequences.

## Introduction

Interferons (IFNs; consisting of types I, II, and III) are secreted cytokines that constitute first line defenses against viral pathogens. They also play key roles in anti-tumor functions as well as in immunomodulatory activities, and as such their importance is emphasized by the evolutionary conservation of IFN genes in all vertebrates (Secombes & Zou, 2017). Classically, IFNs signal by binding to their specific cognate cell surface receptors and triggering activation of Janus kinase (JAK) and signal transducer and activator of transcription (STAT) pathways that lead to the transcription of hundreds of IFN-stimulated genes (ISGs) (Schoggins, 2019). Type I IFNs (IFN-Is), which include 13 subtypes of IFNα as well as IFNβ, IFNω, IFNε and IFNκ, bind to a receptor heterodimer comprised of IFNAR1 and IFNAR2 (Lazear *et al*, 2019; Mesev *et al*, 2019), while the single type II IFN (IFN-II, IFNγ) signals through a receptor composed of IFNGR1 and IFNGR2 subunits (Casanova *et al*, 2024). Type III IFNs (IFN-IIIs) consist of four members, IFNλ1-4, that all share a receptor complex of IFNLR1 and IL10RB (Lazear *et al*., 2019; Mesev *et al*., 2019). Binding of IFNs to their distinct receptor complexes rapidly initiates the kinase activities of their associated cytoplasmic JAK tyrosine kinases (such as JAK1 or TYK2), which primarily act to catalyze well-described phosphorylation events in the STAT proteins (canonically STAT1 or STAT2). These critical tyrosine phosphorylation events drive formation of activated transcription factor complexes in the nucleus: IFN-Is and IFN-IIIs activate the IFN-stimulated gene factor 3 (ISGF3) heterotrimeric complex consisting of phosphorylated STAT1, STAT2 and interferon regulatory factor 9 (IRF9); while IFN-II leads to the formation of a phosphorylated STAT1 homodimer, also known as IFNγ-activated factor (GAF) (Majoros *et al*, 2017). Thus, IFN- triggered tyrosine phosphorylation is essential to propagate the cytokine signal and to ultimately induce ISG-mediated activity (Schoggins, 2019).

Moreover, IFNs have previously been reported to stimulate many additional signaling events in the cell that regulate their actions either positively or negatively. These include those mediated by post-translational modifications such as ubiquitination, SUMOylation, or acetylation, as well as non-tyrosine phosphorylation (Chen *et al*, 2017). For example, in the case of non-tyrosine phosphorylation, type I, II and III IFN stimulation induces serine phosphorylation of STAT1 at S708 and S727, albeit with different kinetics, and both phosphorylation events are required for optimal transcriptional activity (Perwitasari *et al*, 2011; Tenoever *et al*, 2007; Wen *et al*, 1995). In addition, IFNs can activate diverse non- JAK/STAT signaling pathways, such as the PI3K-Akt-mTOR and p38/MAPK cascades, which may collaborate with the JAK/STAT pathway to transmit the IFN signal (Mazewski *et al*, 2020). The activation of the PI3K-Akt-mTOR pathway highlights the role of metabolic and growth-related signaling in the post-transcriptional regulation of immune responses, emphasizing how IFNs can potentially directly influence cellular metabolism and protein synthesis to combat viral infections. In this way, the PI3K-Akt-mTOR axis, while apparently dispensable for ISG transcription, has been implicated in facilitating ISG translation in response to IFN-Is and IFN-II (Kaur *et al*, 2007; Kaur *et al*, 2008; Livingstone *et al*, 2015). The specific signaling pathways activated by IFNs can also vary depending on the cell and IFN type (van Boxel-Dezaire *et al*, 2006). For example, the p38/MAPK pathway has been shown to be essential for IFN-I responses against various viruses, but is not required for IFN-II responses (Li *et al*, 2004). Furthermore, significant crosstalk exists between the IFN signaling cascade and other non-JAK/STAT pathways. A case in point is the non-canonical reprogramming of IFNβ-induced antiviral and inflammatory responses by co-stimulating TNF (Mariani *et al*, 2019). Thus, increasing evidence strongly suggests the cooperation of multiple signaling pathways in the response of cells to different IFN types. Nonetheless, the precise molecular mechanisms underlying such cooperation, as well as some of their functions, remain incompletely understood. Furthermore, it is likely that additional regulatory post-translational modifications triggered by IFNs exist, but have yet to be identified.

In this study, we aimed to characterize global dynamic changes to the human cellular phosphoproteome in response to stimulation with different type I, II and III IFNs. We focused on protein phosphorylation given its reversibility, as well as its known effects on protein conformation, stability, activity, subcellular localization, and interactions (Humphrey *et al*, 2015). We therefore conducted label-free quantitative phosphoproteomics analysis of human lung epithelial (A549) cells treated with IFNα2, IFNβ, IFNω (IFN-I), IFNγ (IFN-II), or IFNλ1 (IFN-III) for various times between 0 and 240 min. Our temporally-acquired data revealed a rich set of > 500 proteins exhibiting differential phosphorylation in response to IFN stimulation, including ∼60 that are commonly regulated by all IFN-Is, and several that exhibit distinct phosphorylation patterns in response to each individual IFN-I, IFN-II or IFN-III. The differentially phosphorylated proteins include examples that have previously been identified to interact with IFN signaling components or whose expression is regulated by IFN. In addition, many differentially phosphorylated proteins and modification sites were identified that have not previously been implicated in IFN function. Using a combination of knock-out and reconstitution assays, siRNA-mediated depletions, gene expression profiling, and biochemical studies, we functionally dissected selected findings from our protein phosphorylation atlas. For example, we reveal the contribution of a previously uncharacterized potential threonine phosphorylation site in STAT1 that regulates IFN-mediated antiviral activity. Furthermore, we highlight a new role for PLEKHG3, a Rho guanine nucleotide exchange factor (RhoGEF) (Bagci *et al*, 2020; Nguyen *et al*, 2016), and its phosphorylation-dependent interaction with 14-3-3 proteins, as collectively fine-tuning IFN signaling. Overall, the specific IFN-regulated phosphoproteomes that we present in this resource will enrich our understanding of host factors and signaling pathways involved in different IFN responses.

## Results

### Widespread Temporal Changes to the Human Proteome and Phosphoproteome During Stimulation with IFNα, IFNβ, IFNω, IFNγ, and IFNλ

To initially explore differences and similarities between the signaling cascades induced by different IFN-Is, we conducted a comparative proteomic and phosphoproteomic survey as outlined in **Fig. 1A**. Human lung epithelial cells, A549s, were independently stimulated with equal amounts of IFNα2, IFNβ, or IFNω (10 ng/ml) for 30, 90 or 240 min. Western blotting confirmed that these conditions similarly activated the canonical JAK/STAT signaling pathway, as demonstrated by a transient increase in STAT1-pY701 levels that peaked at 30 min (**Fig. 1B**). For the proteomic and phosphoproteomic analyses, proteins were extracted post-stimulation from five independent experimental replicates and digested with trypsin, followed by enrichment of the phosphopeptide fraction using Ti^4+^-coupled magnetic beads. Subsequently, peptides were subjected to liquid chromatography tandem-mass spectrometry (LC-MS/MS) with label-free quantification (LFQ). Downstream processing encompassed both total digestions and the phospho-enriched fraction, and peptide identification and LFQ were performed. After conducting quantitative and statistical analyses on the total proteome data, we established a cut-off threshold of two-fold change in average protein abundance between IFN-treated and untreated samples. Following on from this, we calculated the number of proteins exhibiting significantly different abundances (P < 0.01) for each IFN-I and stimulation time (**Fig. 1C**). As expected, the highest number of changes in total protein abundance was observed after 240 min of stimulation with each IFN-I, which (as detailed below) reflects the transcriptional upregulation of ISGs (**Fig. 1C**). Analysis of the IFN-stimulated phosphoproteomes revealed distinct regulatory effects on 238 phosphopeptides for IFNα2, 308 for IFNβ, and 231 for IFNω (**Fig. 1D**). Confirming the known transient nature of IFN signaling and phosphorylation events, most changes in the phosphorylation landscape were detected at the early times after IFN-I stimulation (30 and 90 min) (**Fig. 1D**). Underscoring the validity of our obtained datasets, many of the proteins upregulated after 240 min of stimulation with each IFN-I were indeed well-described ISG protein products, such as MX1, ISG15, TRIM21, OAS1, and IFITs1-3 (**Fig. 1E, Dataset EV1**) (Schoggins, 2019). Similarly, we observed phosphorylation changes in proteins known to be associated with IFN signaling, including STAT1, STAT3, STAT4, and STAT5B (**Fig. 1F, Dataset EV2**) (Hu *et al*, 2021). These results indicate the overall robustness of our methodology, and suggest that the previously unrecognized IFN-regulated proteins and phosphorylation events we have additionally uncovered may play important roles in the response of cells to IFNs.

**Figure 1.**
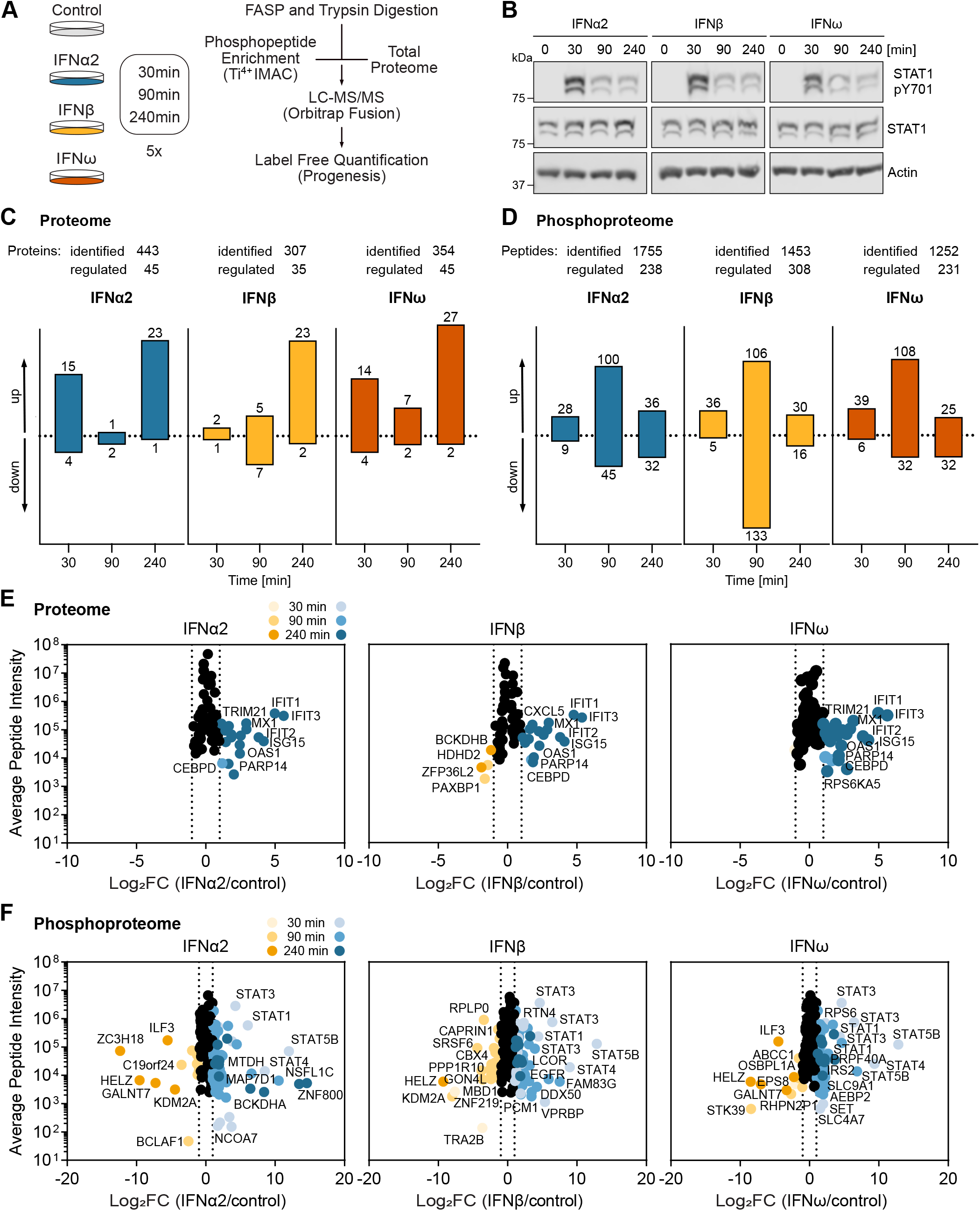
Widespread Temporal Changes to the Human Proteome and Phosphoproteome During Stimulation with IFNα, IFNβ, and IFNω. **A** Schematic overview illustrating the experimental workflow of the phosphoproteomic screen. **B** A549 cells were subjected to overnight serum starvation, followed by stimulation with 10 ng/ml of IFNα2, IFNβ, or IFNω for the indicated durations. Subsequently, cell lysates were analyzed by western blotting with the specified antibodies. Data are representative of n=3 independent experiments. **C-D** Numbers of distinct proteins (**C**) and phosphorylated peptides (**D**) significantly up- or down-regulated (≥ two-fold; P<0.01) at the indicated time points after stimulation with IFNα2 (blue), IFNβ (yellow), or IFNω (red) relative to unstimulated samples. The details of all identified factors are provided in **Dataset EV1** (total proteome) and **Dataset EV2** (phosphoproteome). **E-F** Distribution of changes in the cellular proteome and phosphoproteome in response to IFN stimulation. Log_2_(fold change, FC) in protein abundance (**E**) or phosphorylation (**F**) of the identified peptides of IFN-stimulated samples compared to controls. Horizontal dotted lines represent log_2_(FC) cutoffs of ≤−1 or ≥1. Peptides exhibiting significant changes in abundance are denoted in yellow (decreased) or blue (increased).

To expand our comparisons to include IFN-II and IFN-IIIs, we conducted parallel analyses on A549 cells independently stimulated with IFNγ and IFNλ1. We first confirmed that these two IFNs (at 10 ng/ml) also activated the JAK/STAT signaling pathway in A549s by detecting the stimulated increase in STAT1-pY701 levels using western blot (**Fig. EV1A**). Subsequent proteomic analyses revealed notable and expected changes in total protein abundances following IFNγ or IFNλ1 stimulation, with the most significant alterations observed after 240 min (**Fig. EV1B, Dataset EV1**). Specifically, we observed upregulation of 25 proteins in response to IFNγ and 40 proteins in response to IFNλ1 at this timepoint. Additionally, differential regulation of 160 phosphopeptides was identified upon IFNγ stimulation, and regulation of 356 phosphopeptides was identified in response to IFNλ1 (**Fig. EV1C, Dataset EV2**). Comparison of total proteome changes across all three IFN types (grouping IFNα2, IFNβ, and IFNω together as IFN-I) revealed a subset of 5 proteins that were upregulated in response to all: DTX3L, EPB41L5, ISG15, IFIT3, and SCAMP3; while no overlap was observed for downregulated proteins (**Fig. EV1D-E, Dataset EV3**). Notably, comparative analysis of the IFN-regulated phosphoproteomes indicated a more substantial overlap between IFN types, particularly between IFN-I and IFN-III induced phosphoproteome changes: 57 proteins exhibited a common increase, and 93 a common decrease, in phosphorylation in response to stimulation with these two IFN types (**Fig. EV1F-G, Dataset EV3**). Visualization using Circos plots further highlighted the closer similarities in induced phosphorylation changes among the 3 IFN-Is (IFNα2, IFNβ, and IFNω) as compared to IFN-II and IFN-III (**Fig. EV1H-J**). Overall, this comparative proteomic and phosphoproteomic analysis of the dynamic landscape of IFN-induced signaling has provided a rich resource of common and unique regulated factors that can form the basis for future functional exploration.

### Common and Unique Features of the Phosphoproteomes Regulated by IFN**α**, IFN**β**, IFN**ω**, IFN**γ**, and IFN**λ**

We next focused our attention on assessing the overlap of proteins that change in abundance or phosphorylation upon stimulation with all IFN-Is. We found that 19 proteins increased in expression following stimulation with all three IFN-Is, while zero exhibited a common reduced abundance (**Fig. 2A-B, Dataset EV3**). Notably, many of the common and unique IFN-I-regulated proteins are known ISGs or known interactors of the IFN signaling cascade. Specifically, out of the 85 proteins whose expression changed in response to at least one IFN-I, 13 were previously identified as ISGs (Shaw *et al*, 2017), and 9 are ISGs that interact with at least one of the canonical IFN signaling cascade members: IFNAR1/2, JAK1, TYK2, STAT1/2 or IRF9; according to BioGRID (Oughtred *et al*, 2021) and/or STRING (Szklarczyk *et al*, 2015) (**Fig. 2C, Dataset EV3**). When examining IFN-I-induced phosphoprotein changes, we found that 52 proteins commonly increased in phosphorylation, while 8 proteins commonly decreased in phosphorylation in response to all IFN-Is (**Fig. 2D- E, Dataset EV3**). Furthermore, 40 out of the total 401 proteins that were identified to be phospho-regulated by at least one IFN-I were previously annotated as IFN signaling interactors according to BioGRID (Oughtred *et al*., 2021) and/or STRING (Szklarczyk *et al*., 2015), and 30 proteins were identified as either ISGs, ISGs that also interact with known pathway components, or are associated with IFN signaling according to STRING (Szklarczyk *et al*., 2015) (**Fig. 2F, Dataset EV3**). Considering the timing of total protein abundance changes, all 19 proteins commonly increased by IFN-I stimulation were identified after 240 min of stimulation, consistent with the expected pattern for ISG protein products (**Fig. 2G**). Ten of these ISG proteins were also upregulated by IFNγ, while 6 were also upregulated by IFNλ1 (**Fig. 2G**). In terms of phosphorylation events, we observed changes in 74 distinct phosphopeptides belonging to 56 proteins across all three IFN-Is tested (**Fig. 2H**). Out of these, 21 peptides belonged to proteins previously annotated to interact with IFN signaling members according to BioGRID (Oughtred *et al*., 2021) and/or STRING (Szklarczyk *et al*., 2015), while 4 were recently identified in close proximity to a type I IFN signaling member (Schiefer & Hale, 2024) (**Fig. 2H**). Additionally, 23 phosphopeptides exhibited differential phosphorylation after IFNγ stimulation, and 12 after IFNλ1 stimulation (**Fig. 2H**).

**Figure 2.**
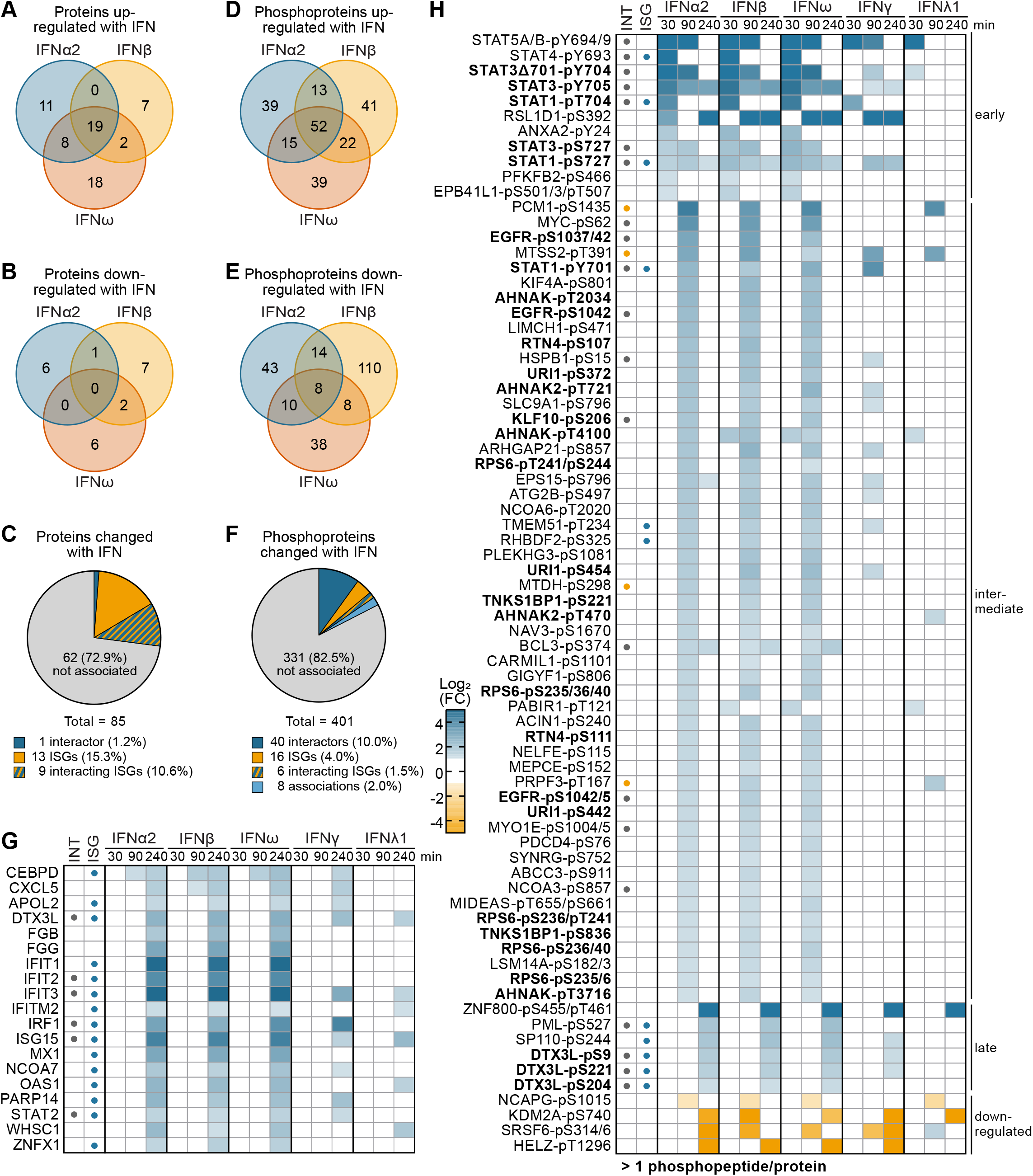
Common and Unique Features of the Phosphoproteomes Regulated by IFN**α**, IFN**β**, IFN**ω**, IFN**γ**, and IFN**λ**. **A-B** Venn diagrams illustrating the overlap in IFN-I-upregulated (**A**) or -downregulated (**B**) proteins following IFNα2, IFNβ, or IFNω stimulation. **C** Pie chart displaying the proportion of IFN-I-changed proteins previously associated with IFN-I signaling according to BioGRID and/or STRING, or considered as ISGs with a more than two-fold increase after IFN stimulation. **D-E** Venn diagrams illustrating the overlap in IFN-I-enhanced (**D**) or -reduced (**E**) phosphoproteins following IFNα2, IFNβ, or IFNω stimulation. **F** Pie chart showing the proportion of IFN-I-changed phosphoproteins previously associated with IFN-I signaling according to BioGRID and/or STRING, or considered as ISGs with a more than two-fold increase after IFN stimulation. The names of all factors in panels **A-F** are provided in **Dataset EV3**. **G-H** Heat maps representing log_2_(fold change, FC) abundances for the 19 proteins (**G**) and for the 74 phospho-peptides (**H**) that change after IFNα2, IFNβ, IFNω, IFNγ or IFNλ1 stimulation. Only log_2_(FC) > 1 and < -1 are depicted. IFN-I signaling interactors (INT) according to BioGRID and/or STRING are indicated with gray dots, and proteins identified in proximity to IFN-I signaling members in a proximity labeling screen (Schiefer & Hale, 2024) are indicated with orange dots. ISGs with a more than two-fold increase after IFN stimulation are indicated with blue dots. Names are written in bold where several distinct phosphorylated peptides were detected for that protein.

Although we observed a strong overlap in phosphoproteomic changes induced by all three IFN-Is, it was clear that each individual IFN-I also triggered its own changes. Specifically, 13 phosphoproteins exhibited unique regulation in response to IFNα2 stimulation, 16 to IFNβ, and 15 to IFNω (**Fig. EV2A-C**). These proteins showed a significant change in phosphorylation (fold change > 2) upon stimulation with only a single IFN, while not being significantly changed with any other tested IFN (irrespective of fold change). In addition, IFNγ stimulation led to 10 proteins undergoing unique phosphorylation changes (**Fig. EV2D**), and no phosphorylation changes were observed to be specific to IFNλ1 alone. While the absence of detectable, significant phosphorylation changes in response to specific IFNs could be a consequence of detection limit or inter-replicate variation in particular experiments, these results may highlight subtle, yet discernible, differences between IFNs in their activated signaling pathways.

To corroborate the regulated phosphorylation events detected by LC-MS/MS, we performed further independent experiments and used western blotting with antibodies specific to selected phosphorylated proteins. For example, we could confirm the increased phosphorylation of STAT1-pS727 and ANXA2-pY24 in response to stimulation with IFNα2 or IFNω (**Fig. EV3A-C**). Moreover, the phosphorylation dynamics observed by this western blot analysis closely resembled the temporal patterns identified by LC-MS/MS (**Fig. EV3D-E**). Overall, these data underscore the reliability and robustness of our experimental approach to elucidate protein phosphorylation dynamics in response to different IFNs. This has revealed both common and unique responses encompassing several factors previously associated with IFN action, as well as many factors that have yet to be functionally implicated in IFN signaling networks.

### Chromatin Remodeling, Transcription, and Splicing are Enriched Biological Processes with Proteins Differentially Phosphorylated Following IFN Stimulation

To gain deeper insights into the functional roles and biological significance of proteins exhibiting differential phosphorylation upon IFN stimulation (particularly IFN-I), we conducted Gene Ontology (GO) enrichment analyses using DAVID (Huang da *et al*, 2009a, b). These analyses revealed enrichment in biological processes such as ’negative and positive regulation of transcription from RNA polymerase II promoter’, ‘mRNA splicing’, and ’chromatin remodeling’ (**Fig. 3A**). Transcriptional regulation mediated by these processes are known to play pivotal roles in modulating gene expression patterns in response to various stimuli, including IFNs (Börold *et al*, 2021; Wang *et al*, 2017). Moreover, examination of the proteins associated with these biological processes revealed a high degree of connectivity (by physical and/or functional association) as depicted in the STRING network analysis (Szklarczyk *et al*., 2015) (**Fig. 3B–D**). Interestingly, proteins involved in the ’positive regulation of transcription from RNA polymerase II promoter’ exhibited predominantly increased phosphorylation levels in response to IFN-I treatment (**Fig. 3B**), whereas those associated with the ’negative regulation of transcription from RNA polymerase II promoter’ tended to show decreased phosphorylation (**Fig. 3C**). The significance of this observation is currently unclear.

**Figure 3.**
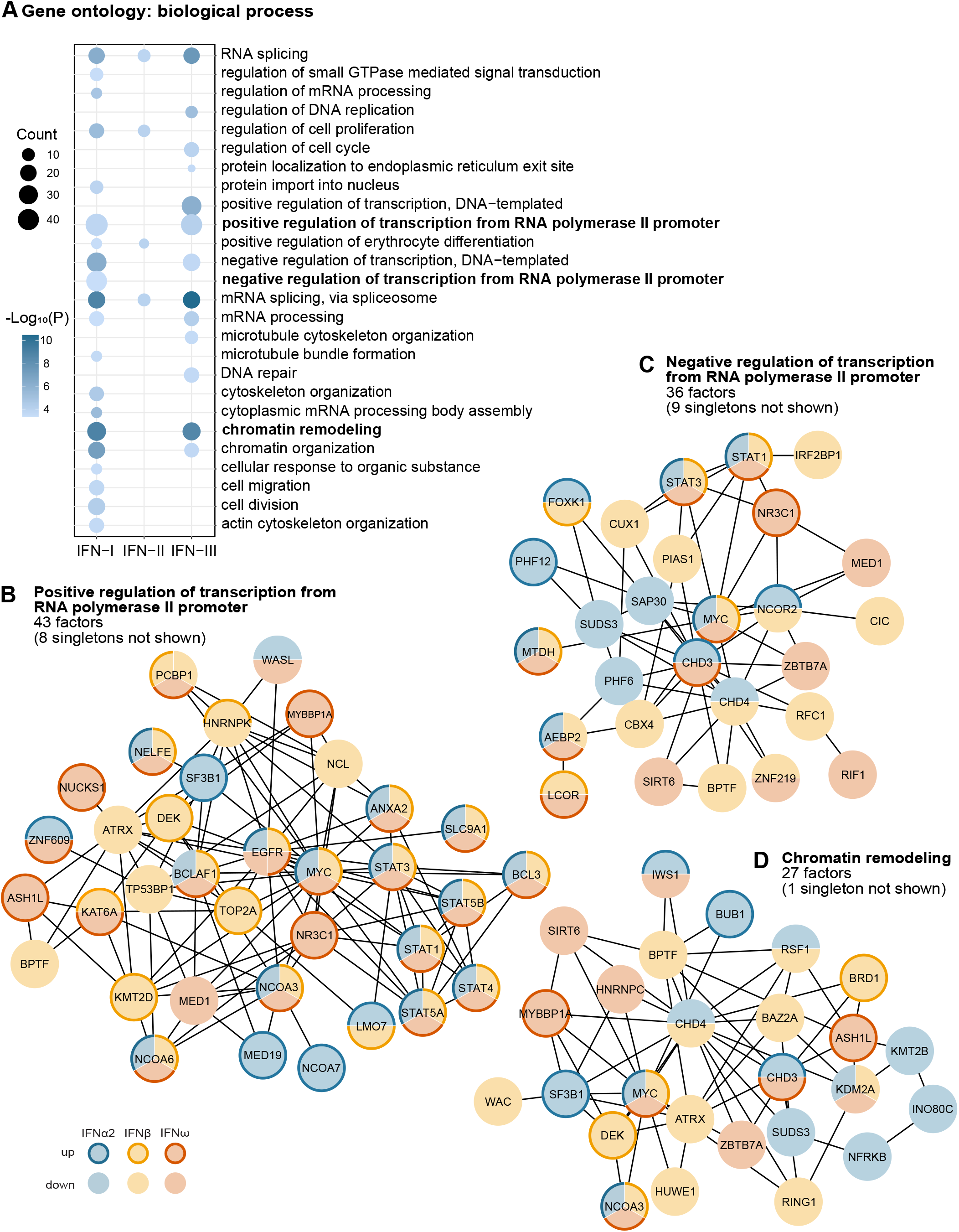
Chromatin Remodeling, Transcription, and Splicing are Enriched Biological Processes with Proteins Differentially Phosphorylated Following IFN Stimulation. **A** Dot plot visualization of enriched GO terms: functional enrichments for biological process gene ontology terms of the identified differentially phosphorylated proteins analyzed using DAVID. Dot color indicates the -log_10_(P) values, while dot size represents the counts of proteins enriched in the GO term. **B-D** Visualization of the STRING network (physical interaction and functional association) of proteins belonging to the significantly enriched biological processes: ’positive regulation of transcription from RNA polymerase II promoter’ (**B**), ’negative regulation of transcription from RNA polymerase II promoter’ (**C**), and ’chromatin remodeling’ (**D**). Unconnected factors (singletons) are not depicted. Networks were generated using Cytoscape 3.9. Factors are highlighted as indicated in the figure key (bold border = increased (up) phosphorylation with IFN; thin border = decreased (down) phosphorylation with IFN).

Not unexpectedly, the GO terms associated with proteins phosphorylated upon treatment with IFN-III (IFNλ1) were similar to those observed with IFN-I (**Fig. 3A**), suggesting a convergence in molecular pathways and biological processes. Both types of IFN influenced the phosphorylation of proteins involved in ‘chromatin remodeling’, ‘RNA splicing’, and ‘positive and negative regulation of transcription’, among others (**Fig. 3A**). In contrast, proteins differentially phosphorylated upon stimulation with IFN-II appeared to be less similar to other IFN types, and there was a markedly diminished number of enriched GO terms that were primarily associated with ‘RNA splicing’ and ‘cell proliferation’ (**Fig. 3A**). It cannot be ruled out that these differences in enriched GO terms between IFN types simply relate to the strength of signaling responses in A549 cells at the doses used.

### STAT1 T704 Contributes to the Full Antiviral Activity of IFN-I

Our phosphoproteomic screen uncovered a previously uncharacterized potential phosphorylation site in STAT1: upon IFN-I stimulation, we observed a phosphopeptide with the well-known phosphorylation of Y701 (90 min post-stimulation), as well as a phosphopeptide assigned with the possible phosphorylation of T704 (30 min post-stimulation) (**Fig. 4A**). Since phosphorylation of STAT1 at T704 has not been previously described, we sought to determine if this residue impacts STAT1 function within the IFN-induced antiviral signaling pathway. To this end, we took advantage of an A549-based STAT1 knockout (KO) cell line (Mazel-Sanchez *et al*, 2021) and used lentiviral vectors to reintroduce wild-type (WT) STAT1 or mutant STAT1s with the amino-acid substitutions T704A or Y701F (as a reference control). Importantly, the lentivirus-derived STAT1 proteins all exhibited expression levels similar to endogenous STAT1 in non-edited control cells (**Fig. 4B, Fig EV4A**), as well as similar STAT1 mRNA levels (**Fig. 4C**). Notably, IFN-stimulated phosphorylation of Y701 could still occur in the STAT1-T704A mutant, and was as efficient as that observed for WT STAT1 when quantified and normalized (**Fig. 4B, Fig EV4B**). Consistent with this observation, stimulation of cells with IFNα2 effectively induced expression of the ISGs, *MX1*, *RIGI* and *IRF9*, in STAT1-KO cells reconstituted with STAT1- WT or -T704A, but not in the non-reconstituted STAT1-KO cells or the KO cells reconstituted with STAT1-Y701F (**Fig 4D-F**). Nevertheless, it was surprising that the cell line expressing STAT1-T704A exhibited reduced IFN-mediated antiviral activity (**Fig. 4G-H**). While pre-treatment of STAT1-WT-expressing cells with IFNα2 led to a significant inhibition of VSV-GFP replication, STAT1-KO cells or cells expressing STAT1-Y701F failed to effectively control the virus following IFNα2 treatment (**Fig. 4G-H**). Strikingly, expression of the STAT1- T704A mutant resulted in an intermediate phenotype, with IFNα2 treatment failing to completely restrict VSV-GFP replication (**Fig. 4G-H**). Similar results were obtained using an IAV-*Renilla* virus (**Fig. 4I**). Interestingly, despite the prior observation that RNA expression levels of selected ISGs were not affected by expression of STAT1-T704A (**Fig 4D-F**), protein levels of these ISGs (most notably MX1) were reduced (**Fig. 4J, K, Fig. EV4C-D**). These data highlight that STAT1 residue T704 may have a role in mediating the antiviral effects of IFN-I independent of global transcription of all ISGs. It is possible that T704 regulates a specific function of STAT1 in promoting ISG translation, at least for some ISGs, but further mechanistic studies will be required to elucidate the precise mechanisms involved, as well as to clarify if phosphorylation at this position is a contributor.

**Figure 4.**
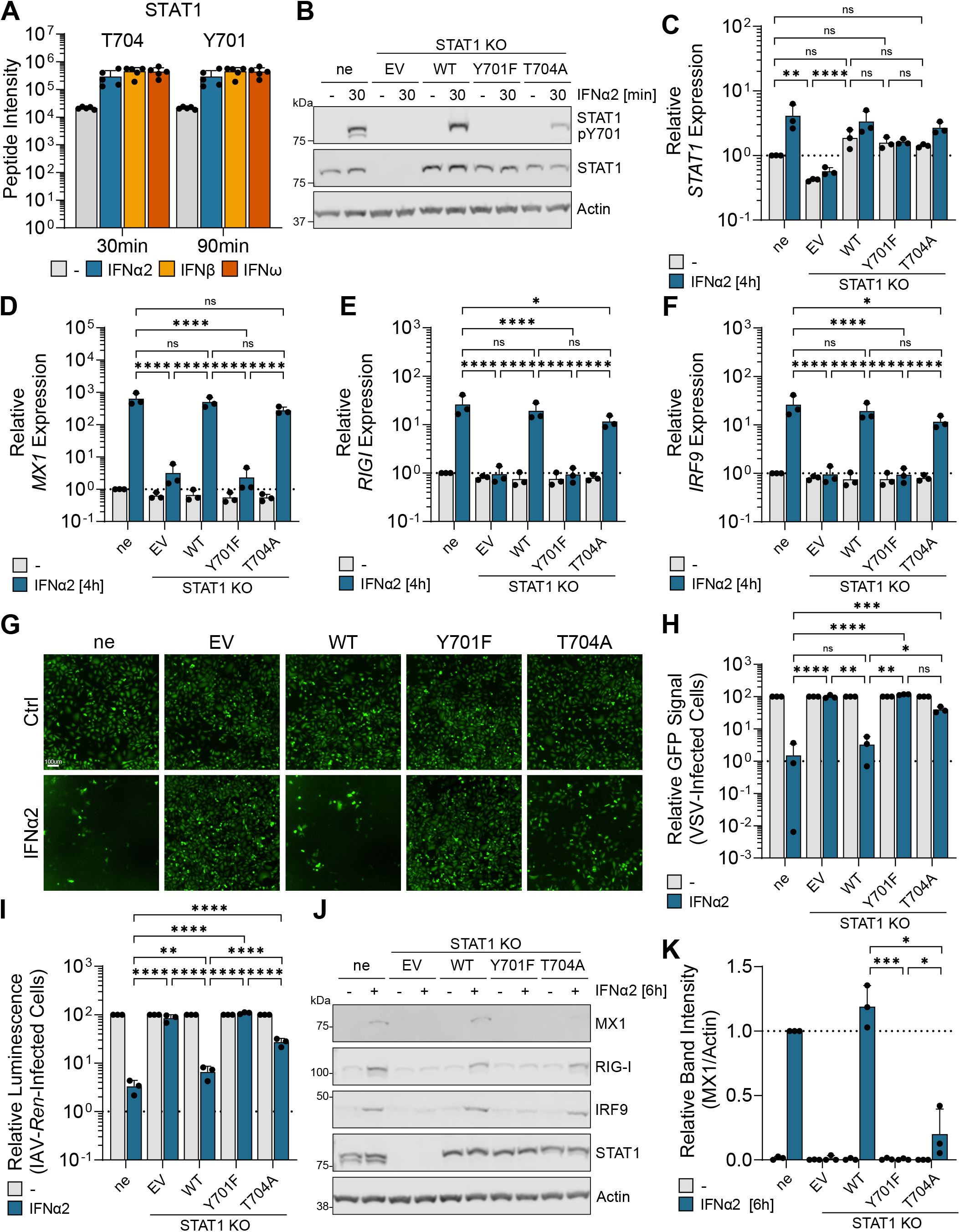
STAT1 T704 Contributes to the Full Antiviral Activity of IFN-I. **A** Phosphopeptide intensities of STAT1 peptides mapped as T704 or Y701 phosphorylation in response to stimulation with each IFN-I. Results from the five independent replicates are shown, and bars represent mean values with standard deviations. **B** Non-edited control A549 cells (ne) or STAT1-deficient (KO) A549 cells reconstituted with empty vector (EV), wild-type (WT) STAT1, or the indicated STAT1 mutants were serum-starved overnight and stimulated with IFNα2 (10 ng/ml) for 30 min. Total protein lysates were analyzed by western blotting with the indicated antibodies. Data are representative of n=3 independent experiments (quantification of three replicates shown in Figure EV4A-B). **C-F** Cells from **B** were stimulated with IFNα2 (10 ng/ml) for 4 h. *STAT1* (**C**), *MX1* (**D**), *RIGI* (**E**) and *IRF9* (**F**) mRNA levels were subsequently determined by RT-qPCR. Relative mRNA expression is shown as fold induction compared to non-stimulated non-edited control cells. **G-H** Cells from **B** were stimulated with 10 ng/ml IFNα2 or not (Ctrl) for 4 h prior to infection with VSV-GFP (MOI = 1 PFU/cell). Representative images (**G**) and quantification of the number of infected cells from n=3 experiments (**H**) are shown for 24 h post infection (hpi). **I** Cells from **B** were stimulated with IFNα2 (10 ng/ml) 16 h prior to infection with *Renilla* luciferase-encoding IAV (WSN/33, MOI = 1 PFU/cell). EnduRen live-cell substrate was added post-infection, and the luciferase activity (relative light units; RLU) was measured at 6 hpi. **J-K** Cells from **B** were serum-starved overnight and stimulated with IFNα2 (10 ng/ml) for 6 h. Total protein lysates were analyzed by western blotting with the indicated antibodies. Data are representative of n=3 independent experiments. Quantification of three replicates is shown for MX1 in (**K**) and for RIG-I, IRF9 and STAT1 in Figure EV4C-E. **C/D/E/F/H/I/K** Bars represent mean values and standard deviations from n=3 independent experiments (each dot corresponds to one replicate). Significance was determined by two-way ANOVA and Tukey’s multiple comparisons test (**C/D/E/F/H/I**) or t-test (**K**) on log-transformed data (*P ≤ 0.05; **P ≤ 0.01; ***P ≤ 0.001; ****P ≤ 0.0001; ns, non-significant).

### RNAi Screening Identifies Novel Regulators of IFN-I Signaling and Antiviral Action Among Phospho-Regulated Proteins

Our proteomic analyses identified many IFN-regulated phosphorylation events in proteins not clearly linked to JAK/STAT signaling. For further characterization, we therefore selected 45 of these proteins that exhibited significant changes in phosphorylation upon stimulation with all three IFN-Is. To assess their functional contribution to IFN-I signaling and antiviral activity, we conducted three different siRNA screening assays based on independent depletion of each identified host factor with four separate siRNAs (**Fig. 5A**). In one set of screens, we assessed the role of each factor in IFN-I-mediated antiviral activity by treating siRNA-transfected cells with 10 ng/ml of IFNα2 for 16 h and then challenging stimulated cells with IAV-*Renilla* or VSV-GFP. We then monitored luminescence (RLU) or GFP expression, respectively, over 12 or 24 h. As a control, we used siRNAs against IRF9, the essential transcription factor for IFN-I signaling, and observed that IRF9 depletion led to a complete inability of IFNα2 to control IAV-*Renilla* or VSV-GFP replication (**Fig. 5A, Dataset EV4**). For our 45 phospho-regulated factors, we considered them potential functional hits when at least 2 out of the 4 siRNAs tested led to a two-fold or greater increase in virus replication following IFNα2 stimulation (summarized in **Fig. 5B, black asterisks in upper and middle panels**). Examples of the full data acquired for the VSV-GFP assay are shown for two resulting functional hits: PML and PLEKHG3 (**Fig. 5C-F**). In a third screen, we transfected siRNAs into an A549 reporter cell line expressing GFP under the control of the *MX1* promoter (A549/pr(ISRE).eGFP cells) (Stewart *et al*, 2014). These cells were then stimulated with 10 ng/ml of IFNα2, and GFP expression was monitored for 72 h using live cell imaging. While positive control siRNAs targeting IRF9 resulted in a clear decrease in IFN-I-induced GFP expression compared to cells transfected with control siRNAs (**Fig. 5A, Dataset EV4**), potential functional hits among our 45 phospho-regulated factors were identified when at least 2 out of the 4 siRNAs tested led to a reduction of >25% in IFN-stimulated GFP expression compared to the control (summarized in **Fig. 5B, black asterisks in bottom panel**). In the three independent screens, we identified 20 host factors that regulate viral replication (14 for VSV-GFP and 12 for IAV-*Renilla*), as well as 15 host factors potentially directly involved in ISG expression (ISRE-GFP assay). Interestingly, several of these candidates have some previous association with antiviral defenses and/or IFN. For example, PDCD4 was shown to play a regulatory role in mRNA translation of ISGs and control of IFN responses by undergoing IFNα-induced phosphorylation and subsequent degradation, facilitating the expression of several important ISG protein products (Kroczynska *et al*, 2012). Furthermore, SRSF6 has been described to limit basal IFN-I expression and apoptosis in macrophages (Wagner *et al*, 2022). Additionally, comparing the results from these three independent screens, we could identify four host factors (PML, PLEKHG3, MTDH, and RSL1D1), whose depletion both enhanced viral replication after IFNα2 stimulation and inhibited IFN-stimulated gene expression. This suggests that these factors may play roles in the IFN signaling response and thus IFN-mediated antiviral activity. Notably, RSL1D1 has recently been identified in a CRISPR/Cas9 knockout screen as a potential regulator of the antiviral host-defense program in response to HSV-1, IAV, and VSV (Pennemann *et al*, 2021). In addition, PML is well-described to possess IFN-regulated antiviral functions (Everett & Chelbi-Alix, 2007). These observations serve as proof of principle that our experimental screening approach may have uncovered new, previously uncharacterized, players in the IFN system from our hit-list of phospho-regulated factors.

**Figure 5.**
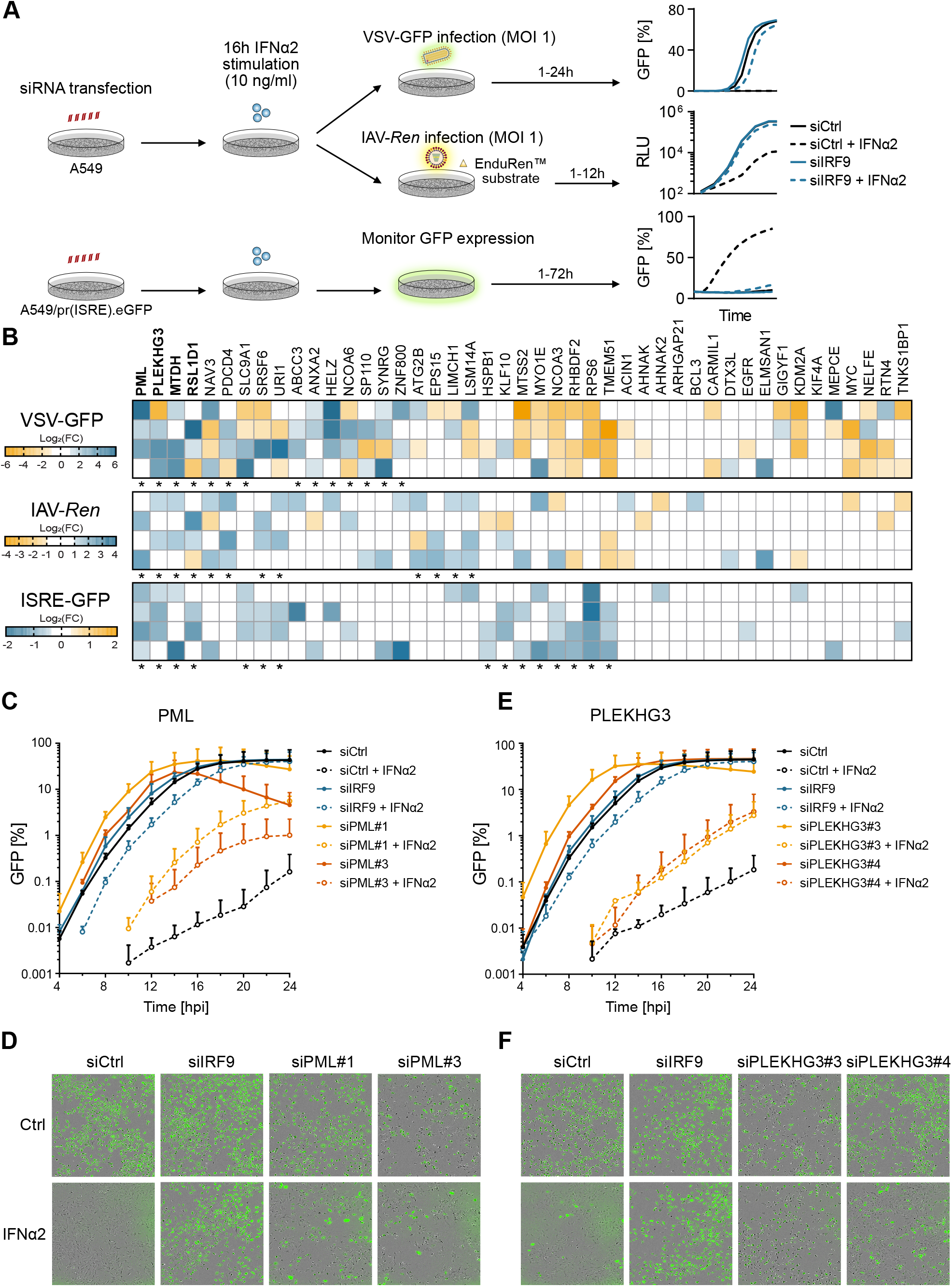
RNAi Screening Identifies Novel Regulators of IFN-I Signaling and Antiviral Action Among Phospho-Regulated Proteins. **A** Schematic representation of the experimental workflows for each RNAi screen. A549 cells or A549 cells expressing GFP under control of the *MX1* promoter (A549/pr(ISRE).eGFP cells) were transfected for 30 h with four independent siRNAs per target, or with a non-targeting control siRNA. Following stimulation with IFNα2 (10 ng/ml) for 16 h, cells were either infected with *Renilla* luciferase-encoding IAV (WSN/33, MOI = 1 PFU/cell) in the presence of the live-cell substrate EnduRen, or with VSV-GFP (MOI = 1 PFU/cell). For IAV- *Renilla* infections of A549 cells, luciferase activity was monitored up to 12 h post infection (hpi), and the area under the curve (AUC) was calculated. For VSV-GFP infections of A549 cells, GFP expression was monitored every 2 h and the percentage of GFP expressing cells was determined. For stimulated A549/pr(ISRE).eGFP cells, GFP expression was monitored every 2 h and the percentage of GFP expressing cells was determined. **B** Heatmaps showing the log_2_(fold change, FC) of GFP or luciferase signals in IFNα2- stimulated siRNA-transfected cells as compared to the non-targeting siRNA control cells. For each gene targeted, the results from the 4 individual siRNAs are shown. Potential hits are indicated with asterisks (at least 2 of the 4 siRNAs led to >2-fold rescue of viral replication in A549 cells, or displayed a >25% reduction in IFNα2-stimulated GFP expression in A549/pr(ISRE).eGFP cells as compared to the relevant control siRNA transfected cells). **C-F** A549 cells were transfected for 30 h with individual siRNAs targeting *PML* (**C-D**), *PLEKHG3* (**E-F**), *IRF9* as a positive control, or a non-targeting siRNA as a negative control. Cells were then stimulated with IFNα2 (10 ng/ml) for 16 h and subsequently infected with VSV-GFP (MOI = 1 PFU/cell). GFP intensity was measured every 2 h. Quantification of the number of infected cells from n=2 independent experiments (**C, E**; means and standard deviations shown) and representative images at 16 hpi (**D, F**) are shown.

### PLEKHG3 has a Role in IFN-I Signaling and Antiviral Activity

From our phospho-regulated factors with screening evidence for functional contributions to IFN-I signaling and antiviral activity, we selected Pleckstrin Homology and RhoGEF Domain Containing G3 (PLEKHG3) for further characterization. PLEKHG3 is a Rho guanine nucleotide exchange factor (RhoGEF) that has not previously been associated with IFN-I (Nguyen *et al*., 2016). We first transfected A549 cells with three independent siRNAs targeting PLEKHG3, and could confirm effective mRNA knockdown by RT-qPCR (**Fig. 6A**). siRNA-transfected cells were also stimulated with varying doses of IFNα2 prior to infection with VSV-GFP and monitoring of infection for 24 h. In line with the initial siRNA screening results, PLEKHG3 depletion could partially rescue VSV-GFP replication from inhibition by IFNα2 stimulation (**Fig. 6B**). Next, we examined if PLEKHG3 knockdown might impact IFN-I signaling by assessing ISG transcription following IFNα2 stimulation. Indeed, knockdown of PLEKHG3 with all three independent siRNAs significantly reduced IFNα2-induced expression of several ISGs (*MX1, IFIT1, IFIT2, RIGI, IFITM3, RSAD2*) (**Fig. 6C-E and Fig. EV5A-C**). To determine if this effect was specific to signaling downstream of stimulation with IFNα2, we next depleted PLEKHG3 using an siRNA pool (**Fig. 6F**), and stimulated cells with various IFN-Is (IFNα2, IFNβ, IFNω), IFN-II (IFNγ), and IFN-III (IFNλ1), before assessing ISG expression (**Fig. 6G-H**). While stimulation with all these IFNs led to detectably enhanced expression of *MX1* and *IFIT2*, the effect of PLEKHG3 knockdown on limiting IFN-stimulated gene expression was specific to the IFN-Is and IFN-III, and had no significant effect on IFN-II-induced ISGs (**Fig. 6G-H**). This specificity was further confirmed by assessing TNFα signaling, where PLEKHG3 depletion appeared to increase TNFα-stimulated *NFKBIA* expression (**Fig. 6I**). Notably, PLEKHG3 depletion did not affect early events in the IFN-I signaling cascade, such as IFN-I-stimulated STAT1 or STAT2 phosphorylation (**Fig. EV5D**).

**Figure 6.**
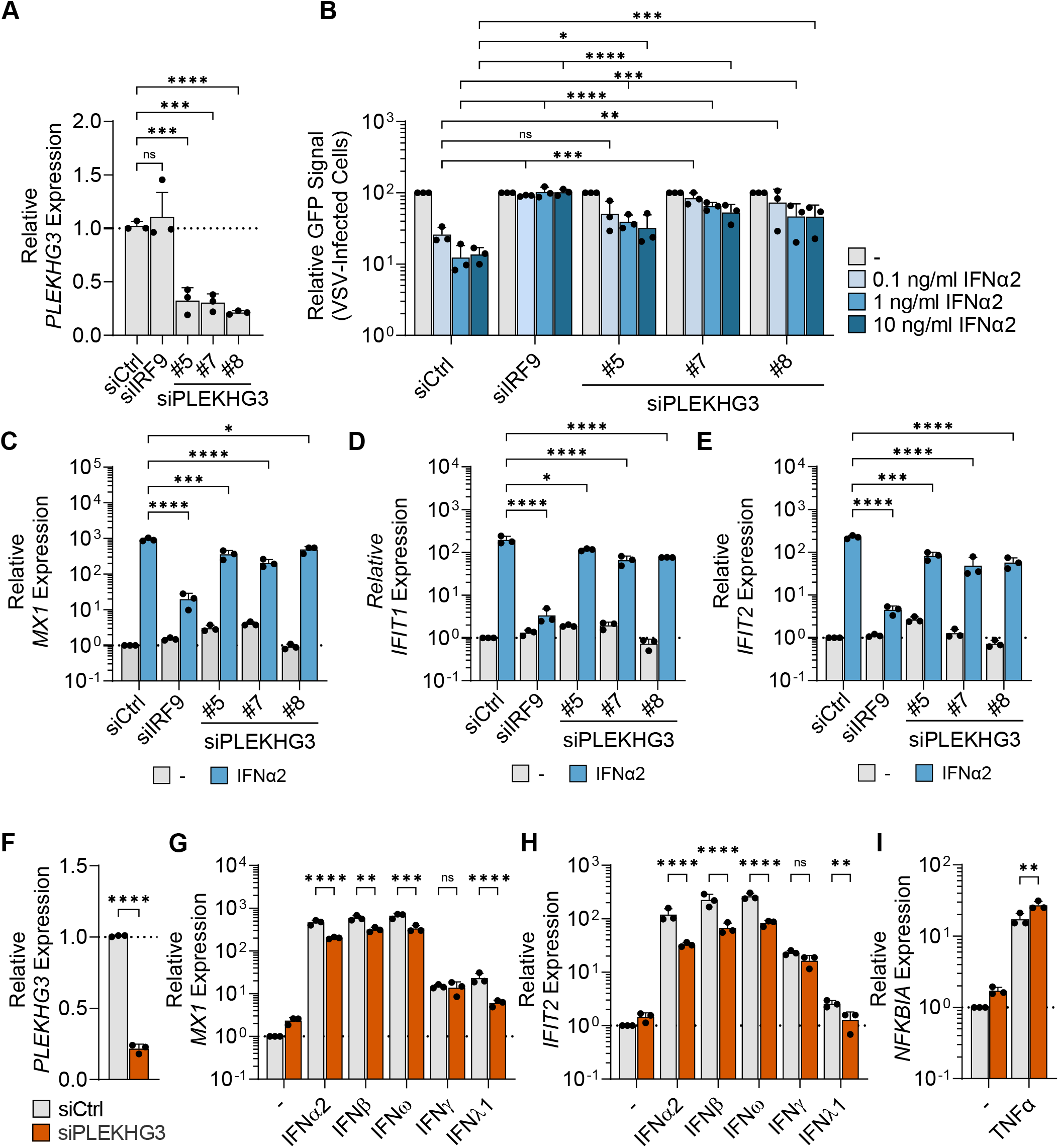
PLEKHG3 has a Role in IFN-I Signaling and Antiviral Activity. **A** A549 cells were transfected for 30 h with individual siRNAs targeting *PLEKHG3*, *IRF9* as a positive control, or a non-targeting siRNA as a negative control. Total RNA was harvested, and expression of *PLEKHG3* mRNA was assessed by RT-qPCR using specific primers. **B** A549 cells were transfected as in **A**, before stimulation with different concentrations of IFNα2 for 16 h. Cells were then infected with VSV-GFP (MOI 1 PFU/cell) and GFP expression assessed at 24hpi. **C-E** A549 cells were transfected as in **A**, before stimulation with IFNα2 for 4 h (1 ng/ml). Total RNA was harvested, and expression of the indicated genes was assessed by RT-qPCR using specific primers. **F-I** A549 cells were transfected for 30 h with a pool of 3 siRNAs targeting *PLEKHG3* or with a non-targeting siRNA as a negative control. Cells were then stimulated with different IFNs for 4 h (1 ng/ml IFNα2, 0.12 ng/ml IFNβ, 1 ng/ml IFNω, 5 ng/ml IFNγ, or 5 μg/ml IFNλ1) (**G/H**) or TNFα for 1 h (10 ng/ml) (**I**). Total RNA was harvested, and expression of the indicated genes was assessed by RT-qPCR using specific primers. **A-I** Bars represent mean values and standard deviations from n=3 independent experiments (each dot corresponds to one replicate). Significance was determined by ordinary one-way ANOVA with Dunnett’s multiple comparison test (**A**), two-way ANOVA with Dunnett’s multiple comparison test (**B-E**), unpaired t-test (**F**), or two-way ANOVA with Šídák’s multiple comparisons test (**G-I**) on log-transformed data (*P ≤ 0.05; **P ≤ 0.01; ***P ≤ 0.001; ****P ≤ 0.0001; ns, non-significant).

These data indicate that PLEKHG3 contributes specifically to IFN-I/III-stimulated gene expression and antiviral activity, but this function is independent of early steps in the IFN signaling cascade.

### Phosphorylation of PLEKHG3 at S1081 can be Mediated by an IFN-Regulated Kinase and is Required for Interaction with 14-3-3 Proteins

Our initial phosphoproteomic analysis revealed that PLEKHG3 is phosphorylated at S1081 in response to stimulation with IFN-Is after 90 min (**Fig. 7A**). While examining the sequence of PLEKHG3 surrounding the S1081 phosphorylation site, we noticed that it resembles a 14-3- 3 consensus binding motif (**Fig. 7B**) (Liu *et al*, 2021). 14-3-3 proteins are evolutionarily conserved scaffolding molecules that modulate the function of other proteins through phosphorylation-dependent interactions, regulating processes like signal transduction, apoptosis, autophagy, and cell cycle regulation (Liu *et al*., 2021; Obsilova & Obsil, 2022). Mammals express seven different 14-3-3 proteins (β, γ, ε, ζ, η, θ, σ), and some 14-3-3 proteins have already been implicated in regulating IFN-mediated antiviral defenses, for example by interacting with RIG-I and MDA5 (Lin *et al*, 2019; Liu *et al*, 2012), or by binding to STAT3 (Munier *et al*, 2021). We verified the presence of the 14-3-3 binding motif in PLEKHG3 by using an antibody that specifically recognizes the phosphorylated 14-3-3 binding motif in target proteins. Flag-tagged PLEKHG3 ectopically expressed in HEK293T cells could be readily recognized by this antibody via western blot, and this detection was sensitive to λ-phosphatase treatment revealing the specificity to phosphorylated PLEKHG3 (**Fig. 7C**). Furthermore, a PLEKHG3 mutant in which the S1081 phosphorylation site was substituted to alanine (S1081A) was no longer detected by the 14-3-3 phospho-binding motif antibody (**Fig. 7C**). Among the few serine kinases reported to contribute to IFN signaling, IKKε is known to phosphorylate STAT1 at S708 in response to IFNβ stimulation (Perwitasari *et al*., 2011). Thus, we tested whether IKKε might also mediate PLEKHG3 phosphorylation. Therefore, ectopically expressed PLEKHG3 was immunoprecipitated from HEK293T cells, dephosphorylated with λ-phosphatase, and subjected to an *in vitro* kinase assay with recombinant IKKε. The results confirmed that IKKε is indeed capable of phosphorylating PLEKHG3 *in vitro* (**Fig. 7D**). Next, we set out to analyze the interaction of ectopically expressed PLEKHG3 with each of the seven 14-3-3 family members by co-immunoprecipitation. PLEKHG3 interacted with all seven 14-3-3 proteins, but exhibited a slight preference for 14-3-3γ and 14-3-3θ (**Fig. 7E-F**). The interaction between PLEKHG3 and 14-3-3θ required the intact 14-3-3 binding motif in PLEKHG3, and in particular the S1081 phosphorylation site, as the PLEKHG3-S1081A mutant did not interact with 14-3-3θ (**Fig. 7G**). These findings indicate that the motif surrounding PLEKHG3-S1081 is a *bona fide* functional 14-3-3 binding motif.

**Figure 7.**
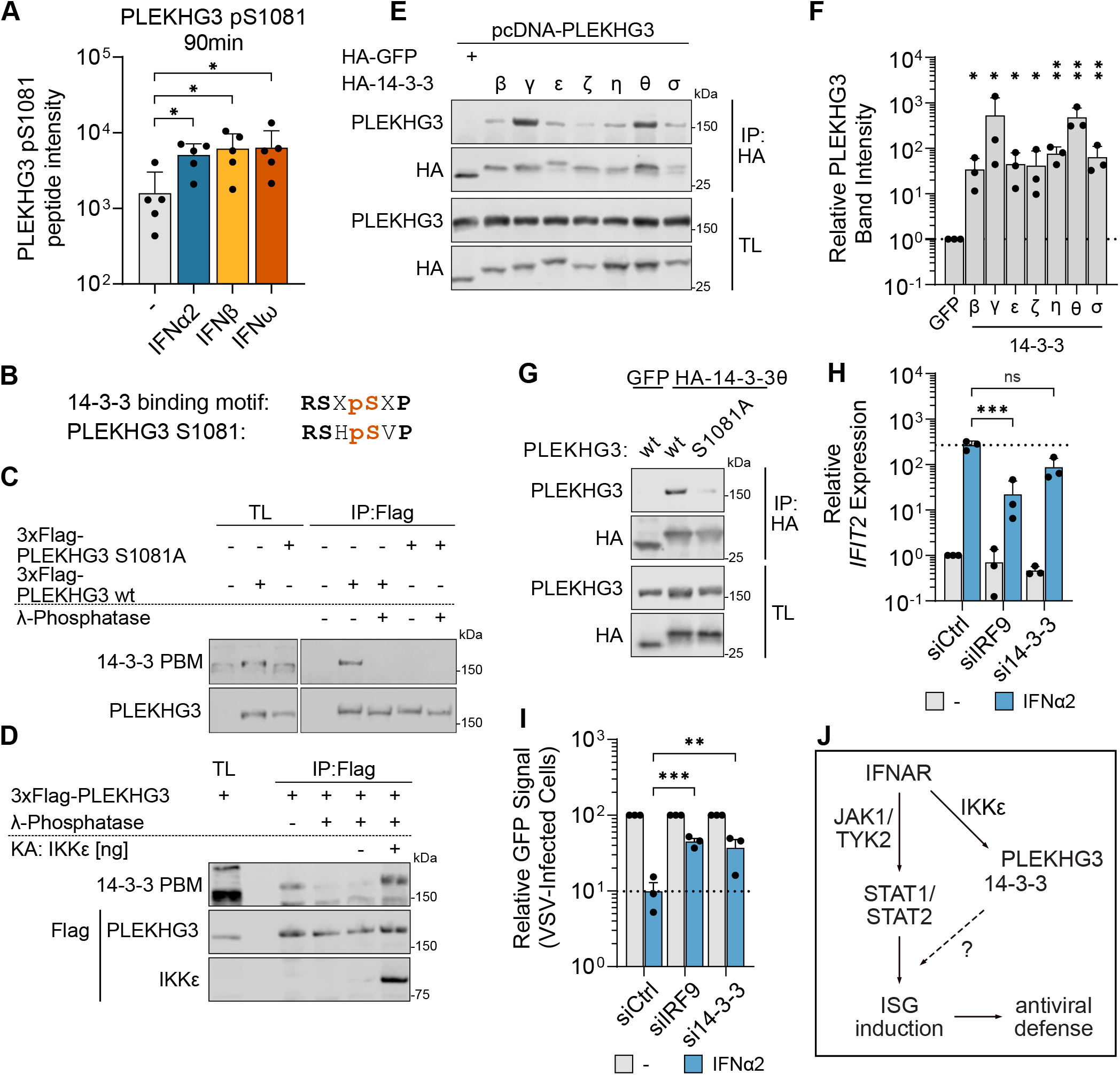
Phosphorylation of PLEKHG3 at S1081 can be Mediated by an IFN-Regulated Kinase and is Required for Interaction with 14-3-3 Proteins. **A** Phosphopeptide intensities of the PLEKHG3 pS1081 phosphopeptide after stimulation with IFNα2, IFNβ or IFNω for 90 min. Results from the five independent replicates are shown, and bars represent mean values with standard deviations. Significance was determined by one-way ANOVA (*P ≤ 0.05). **B** Depiction of a consensus 14-3-3 binding motif and the corresponding sequence in PLEKHG3 surrounding the phosphorylation site at S1081. pS represents phosphorylated serine, and X represents any amino-acid. **C-D** HEK293T cells were transiently transfected with 3xFlag-PLEKHG3 wt or the S1081A mutant. PLEKHG3 was then immunoprecipitated from lysates using an antibody against the Flag tag. Immunocomplexes were treated with λ-phosphatase where indicated, and subjected to an *in vitro* kinase assay (KA) with recombinant Flag-IKKε (**D**). Immunoprecipitates (IP) and total lysates (TL) were analyzed by western blotting with the indicated antibodies. Data shown are representative of at least n=3 independent experiments. **E-G** HEK293T cells were transiently transfected with 3xFlag-PLEKHG3 wt or the S1081A mutant, together with HA-tagged 14-3-3 proteins or GFP as indicated. 14-3-3 proteins or GFP were immunoprecipitated from lysates using an antibody against the HA-tag. Immunoprecipitates (IP) and total lysates (TL) were analyzed by western blotting with the indicated antibodies. PLEKHG3 band intensities in immunoprecipitates (IP) in **E** were quantified using Image Studio Lite and are shown in **F**, where bars represent mean values and standard deviations from n=3 independent experiments (each dot corresponds to one replicate). Significance was determined by one-sample t-test on log-transformed data (*P ≤ 0.05; **P ≤ 0.01). **H-I** A549 cells were transfected with negative-control siRNAs, an siRNA targeting *IRF9*, or siRNA pools targeting all seven 14-3-3 family members (14-3-3β, 14-3-3γ, 14-3-3ε, 14-3-3ζ, 14-3-3η, 14-3-3θ, 14-3-3σ). 48 h post transfection, cells were stimulated with IFNα2 (1 ng/ml) for 4 h. Some cells were harvested and *IFIT2* mRNA levels were determined by RT- qPCR (**H**). Other cells were infected with VSV-GFP (MOI = 1 PFU/cell) and GFP expression as a measure for viral replication was assessed at 24hpi (**I**). Bars represent mean values and standard deviations from n=3 independent experiments (each dot corresponds to one replicate). Significance was determined by two-way ANOVA with Dunnett’s multiple comparison test on log-transformed data (**P ≤ 0.01; ***P ≤ 0.001; ns, non-significant). **J** Schematic model for the role of the PLEKHG3:14-3-3 axis in IFN signaling: we suggest that IFN-activated IKKε phosphorylates PLEKHG3 and facilitates the interaction of PLEKHG3 with one or more 14-3-3 proteins to regulate signal transduction downstream of STAT1/STAT2 phosphorylation.

Our identification of PLEKHG3-S1081 phosphorylation following IFN stimulation (potentially via the IFN-regulated kinase IKKε) and its subsequent recognition by a broad set of 14-3-3 proteins led us to next characterize the role of 14-3-3 proteins in IFN signaling. Thus, we used an siRNA-mediated knockdown approach to independently target all the 14-3-3s in A549 cells, and analyzed the impact of this on IFNα2-induced ISG expression and antiviral activity. Surprisingly, knockdown of individual 14-3-3s either increased or decreased ISG induction depending on the target, and knockdown also either enhanced or suppressed the antiviral activity of IFN-I (**Fig. EV6A-B**). For example, depletion of 14-3-3η enhanced IFN-stimulated *IFIT2* expression, while knockdown of 14-3-3γ reduced it. Conversely, VSV-GFP replication in the presence of IFN-I was partially rescued in cells depleted of 14-3-3σ, but was more strongly inhibited when 14-3-3η was depleted. These data suggest that there are distinct and complex contributions of different 14-3-3 proteins towards IFN signaling and antiviral activity, which is consistent with their wide-ranging roles as scaffolding molecules. Interestingly, collective knockdown of all seven 14-3-3s partially reduced IFNα2-stimulated *IFIT2* expression (although this was not statistically significant), and significantly limited the antiviral effect of IFNα2 (**Fig. 7H-I**). These data underscore the overall positive role that 14- 3-3 proteins have in IFN-mediated antiviral activity. Given the positive role that PLEKHG3 also has in this pathway, the IFN-induced phosphorylation of PLEKHG3 at S1081, potentially by IKKε, may create a 14-3-3 docking platform for these factors to collectively regulate IFN signaling and antiviral responses.

## Discussion

Herein, we employed an unbiased proteomics approach to identify similarities and differences between several IFNs (α2, β, ω, γ, and λ1) with respect to their temporal regulation of cellular phosphorylation events. This led to the identification of >700 total IFN-regulated phosphorylation sites in >500 proteins, vastly increasing our current understanding of factors associated with these pathways. The results we present differentiate themselves from other phosphoproteomic efforts in this area (for example (Vázquez-Blomquist *et al*, 2022; Viengkhou *et al*, 2021)) due to the short IFN stimulation periods that we used to capture rapid and transient events relevant to the known timings of JAK/STAT signaling, as well as the comprehensive nature of our approach using all three IFN types. Furthermore, the work adds a complementary additional layer of knowledge to recent studies that have investigated the common and unique specificities of different IFNs with regards to their antiviral activities, or their transcriptome and proteome re-wiring capabilities (Guo *et al*, 2020; Lum *et al*, 2024; Matos *et al*, 2019; Megger *et al*, 2017; Schuhenn *et al*, 2022; Yan *et al*, 2004). Thus, our datasets reported here cover a breadth of IFN-induced phosphorylation changes in multiple cellular pathways and should be a valuable additional resource that supports the community in undertaking future studies to dissect the actions and consequences of IFN stimulation.

Our own follow-up studies focused on the characterization of IFN-regulated phosphorylation events in both known and previously undescribed components of the IFN signaling pathway, with a particular focus on common IFN-I-induced changes. One such finding was the potential observation of a previously undescribed potential phosphorylation event in STAT1 at T704, close to the critical Y701 residue. Using a knock-out and reconstitution strategy to express wild-type and mutant STAT1 proteins, we found that T704 plays an important role in mediating the antiviral activity of IFN-I, but in a manner seemingly independent of regulating global ISG transcription. In fact, T704 in STAT1 appears to support a later step in mediating the antiviral activity of IFN-I, possibly at the level of translation of selected ISGs, as substitution of T704 in STAT1 compromised IFN-induced levels of some ISG proteins, but not mRNAs. It is intriguing to speculate that IFN-I may trigger phosphorylation of T704 to regulate the actions of STAT1 signaling, particularly in the context of differentiating a role for STAT1 in transcription versus translation. In this context, it is interesting to note that STAT1 has previously been described to promote cap-independent mRNA translation of selected mRNAs in cancer cell proliferation (Wang *et al*, 2015). STAT1 T704 phosphorylation remains to be confirmed, but if validated it is possible that this phosphorylation may affect its interaction with other effector molecules. For example, it is notable that T704 is adjacent to a known SUMOylation site in STAT1, K703 (Rogers *et al*, 2003). SUMOylation at this specific site impedes STAT1 nuclear translocation and its ability to bind DNA (Begitt *et al*, 2011; Grönholm *et al*, 2012; Ungureanu *et al*, 2005). Additionally, it has been demonstrated that K703-SUMOylation is mutually exclusive with Y701-phosphorylation (Zimnik *et al*, 2009). Consequently, it can be hypothesized that potential phosphorylation at T704 may regulate STAT1 SUMOylation, thereby redirecting its mode of action.

Given that our IFN-I-induced phosphorylation profiling identified post-translational changes to many factors not previously described in the canonical JAK/STAT pathway, we also sought to understand if any of these also contribute to the antiviral IFN-I response. Thus, our use of three complementary independent loss- and gain-of-function RNAi screens revealed several potential regulators of the IFN signaling cascade and/or antiviral activity among the proteins differentially-phosphorylated in response to IFN-I. Notable factors identified in these screens included PML, PLEKHG3, MTDH, RSL1D1, SLC9A1, SRSF6, and URI1 that, when knocked-down, all reduced IFN-I-mediated ISG induction and increased virus replication, indicating a positive role for these factors in IFN-I signaling. One of those factors, PML, is an ISG that has previously been shown to positively regulate IFN-II signaling (El Bougrini *et al*, 2011) and to be required for IFN-I-mediated induction of apoptosis (Crowder *et al*, 2005; Herzer *et al*, 2009). In addition, several other factors in our screens appeared to play positive roles in mediating the antiviral activity of IFN-I without appreciably impacting IFN-I-induced *MX1* promoter based expression: NAV3, PDCD4, ABCC3, ANXA2, HELZ, NCOA6, SP110, SYNRG, ZNF800, ATG2B, EPS15, LIMCH1, and LSM14A. Among these, PDCD4 is known to play a regulatory role in translation of ISGs (Kroczynska *et al*., 2012), SP110 is an ISG with important antiviral effects (Fraschilla & Jeffrey, 2020; Shaw *et al*., 2017), ATG2B is involved in autophagy and potentially immune-regulation (Liu *et al*, 2023), and LSM14A is a sensor of viral infections (Li *et al*, 2012). While for many of these factors the precise role that phosphorylation may have on their function is currently unknown, our work presented here identifying their IFN-I-regulated post-translational modification and impact on IFN-I-regulated antiviral activity provides a critical phenotypic framework to explore this in the future.

We characterized PLEKHG3, and its IFN-I-regulated phosphorylation, in more detail. PLEKHG3 is one of eight members of the Pleckstrin homology domain-containing family G (PLEKHG) proteins, and is a RhoGEF that accumulates in focal adhesions through its binding to actin (Nguyen *et al*., 2016). It has been described to affect lysosomal trafficking and cell motility (Ettelt *et al*, 2023), but to our knowledge there are no previous reports connecting PLEKHG3 with the IFN response. We show here that PLEKHG3 is specifically important for gene expression in response to IFN-I and IFN-IIIs, but not to IFN-II or signaling downstream of TNFα. Furthermore, we demonstrate that the IFN-I-promoted phosphorylation of PLEKHG3 at residue S1081 can be performed by IKKε (a known IFN-regulated kinase (Tenoever *et al*., 2007)) and creates a docking motif for interaction with many different 14-3-3 proteins, which, *in globo* at least, we show are also important for IFN signaling and antiviral responses. 14-3-3 proteins have previously been implicated in regulating immune and inflammatory responses (Munier *et al*., 2021), and 14-3-3 proteins may collectively act as a rheostat for intracellular signaling in general, buffering acute phosphorylation outbursts (Gogl *et al*, 2021). However, the molecular mechanisms by which 14-3-3 proteins regulate their binding partners’ functions are still not fully understood (Obsilova & Obsil, 2022), and our own mechanistic characterization of the phospho- PLEKHG3:14-3-3 axis was complicated by the observation that depletion of individual 14-3-3 proteins variably affected the IFN pathway: knockdown of some 14-3-3s tended to enhance IFN-I-induced ISG expression and antiviral activity, while depletion of other 14-3-3s tended to reduce IFN-I-induced ISG expression and antiviral activity. Other studies have also shown that specific 14-3-3 proteins can impact JAK/STAT function: 14-3-3ζ interacts with STAT3 to stabilize its S727 phosphorylation and thereby positively regulate downstream signaling (Zhang *et al*, 2012), and it may be that other 14-3-3 proteins act in an antagonistic manner. Taken together with our findings, we therefore suggest that IFN-activated IKKε phosphorylates PLEKHG3 and facilitates the interaction of PLEKHG3 with one or more 14-3- 3 proteins (**Fig. 7J**). This process likely occurs simultaneously with, or shortly after, IFN-mediated JAK/TYK activation, as phosphorylation of PLEKHG3-S1081 was observed 90 minutes post-IFN-I stimulation. Furthermore, PLEKHG3 depletion did not affect JAK-mediated tyrosine phosphorylation of STAT1/STAT2, suggesting that the PLEKHG3:14-3-3 axis does not play a role in regulating this process. Rather, we speculate that the phosphorylation of PLEKHG3 could either act to sequester specific ‘negative-regulator’ 14-3- 3s away from STAT signaling components, or to aid delivery of other specific ‘positive-regulator’ 14-3-3s to STATs. In this latter regard, the 14-3-3s could influence the heterocomplex formation, dimerization, nuclear translocation, or sustained activation of STAT1/STAT2 signaling, similar to the reported regulation of STAT3 by 14-3-3ζ (Zhang *et al*., 2012). We note that PLEKHG3 depletion only affected IFN-I and IFN-III (but not IFN-II) signaling, therefore it is intriguing to further speculate that the component targeted by such a mechanism would be STAT2, the key factor unique to these pathways.

In summary, our atlas of protein phosphorylation temporal dynamics triggered by type I, II and III IFN stimulation is a rich resource with which to further study the complexities of IFN-mediated antiviral and antiproliferative activity. As examples herein, we could begin to functionally dissect the antiviral contribution of previously unappreciated phosphorylation events in novel factors. Future comprehensive characterization of the IFN-regulated protein phosphorylation landscape will no doubt provide valuable insights into the interplay between IFN responses and a wide array of cellular processes. Indeed, while IFNs have been promising candidates as therapeutic agents against some viral infections, cancers, and autoimmune malignancies, there are poorly understood side-effects elicited by this treatment regimen (De Ceuninck *et al*, 2021; Lasfar *et al*, 2016a; Lasfar *et al*, 2016b; Li *et al*, 2018; Shen *et al*, 2018). Thus, beyond a basic understanding of cellular antiviral mechanisms, our work may help to uncover the complex molecular consequences of IFN stimulation across multiple pathways. This could support a better understanding of both desirable and non-desirable IFN-regulatory effects, and may therefore allow improved rational design of cytokine-based therapeutic strategies.

## Material and Methods

### Cells, viruses, and proteins

A549 and HEK293T cells were cultured in Dulbecco’s modified Eagle’s medium (DMEM) (Life Technologies) supplemented with 10% (v/v) FBS, 100 U/ml penicillin, and 100 μg/ml streptomycin (Gibco Life Technologies). A549/pr(ISRE).eGFP cells were a kind gift from Rick Randall and Catherine Adamson (University of St Andrews, UK) (Stewart *et al*., 2014). A549-STAT1-KO cells have been described previously (Mazel-Sanchez *et al*., 2021), and for reconstitution experiments were transduced for 48 h with the respective lentiviral particles (see below) in the presence of polybrene (8 μg/ml, Sigma-Aldrich) prior to selection with puromycin (1 μg/ml; Thermo Fisher Scientific). VSV-GFP was a kind gift from Peter Palese (Icahn School of Medicine, New York, USA) and IAV-*Renilla* (WSN/33 strain) has been described previously (Spieler *et al*, 2020). The following compounds were used for stimulations: IFNα2 (Novusbio, NBP2- 34971) IFNβ1b (Novusbio, NBP2-35892), IFNω (Novusbio, NBP2-35893), IFNγ (Novusbio, NBP2-34992), IFNλ1 (Novusbio, NBP2-34996) and recombinant human TNFα (PeproTech, 300-01A).

### Transfections and plasmids

A549 cells were transiently transfected with siRNAs using Lipofectamine RNAiMAX transfection reagent (Thermo Fisher Scientific) according to the manufacturer’s instructions. The target sequences of siRNAs used are listed in **Dataset EV5**. Cells were used for downstream applications 48 h post transfection. For transient transfection of plasmids, 7.5x10^5^ HEK293T cells were seeded per well of a 6-well plate, and transfected the next day using FuGene HD transfection reagent (Promega) at a 1:3 DNA:transfection reagent ratio. For lentiviral production, HEK293T cells were transfected as indicated above with an appropriate pLVX-IRES-Puro plasmid, together with psPAX2 and pMD2.G (Addgene plasmids #12259 and #12260; gifts from Didier Trono) at a ratio of 2:1:1. At 48 h post-transfection, supernatants were harvested and filtered through a 0.45 µm syringe filter.

The following additional plasmids were used in this study: pMK-PLEKHG3 was generated by Invitrogen GeneArt Gene Synthesis (Thermo Fisher Scientific) and the PLEKHG3 cDNA was sub-cloned into p3xFlag and pcDNA3.1 vectors using EcoRI and XbaI. pCS2-HA-14-3-3ε (Addgene #116886), pCS2-HA-14-3-3ζ (Addgene #116888) and pCS2-HA-14-3-3η (Addgene #116887) were a gift from Feng-Qian Li & Ken-Ichi Takemaru (Li *et al*, 2008); pEMB52-14-3-3_1_247 (γ) was a gift from the Michael J Fox Foundation MJFF (Addgene #40541); pcDNA-HA-14-3-3β (Addgene #13270), pcDNA-HA-14-3-3σ (Addgene #11946) and pGEX-4T2-14-3-3 tau (θ) (Addgene #13281) were gifts from Michael Yaffe (Yaffe *et al*, 1997). Where necessary, 14-3-3 cDNAs were sub-cloned into pCS2-HA using EcoRI and XhoI so that they could all be expressed with an N-terminal HA-tag. EGFP cDNA was similarly sub-cloned from an existing vector, pLVX-EGFP (Fernbach *et al*, 2022), into pCS2- HA using EcoRI and XhoI. STAT1 cDNA was PCR-amplified from Addgene #8691 (a gift from Jim Darnell (Horvath *et al*, 1995)) and sub-cloned into pLVX-IRES-Puro (Takara) via the BamHI and NotI sites. Indicated mutations were introduced with the QuikChange Site-Directed Mutagenesis kit (Agilent) using the primers listed in **Dataset EV5**.

### Proteomic sample preparation and analysis

A549 cells were serum-starved for 16 h before stimulation with 10 ng/ml of IFNα2, IFNβ, IFNω, IFNγ or IFNλ1. After 30, 90, or 240 min, cells were lysed in SDS-lysis buffer (4% (w/v) SDS, 100mM Tris/HCL pH 8.2, 0.1M DTT, Complete (Roche) and Phos-Stop (Roche)) followed by a short sonication to reduce viscosity. Protein amounts were then measured using a Qubit fluorometer (Invitrogen). Five independent experimental replicates were performed. Tryptic digests and phosphopeptide enrichment were performed as described in previous detailed protocols (Hunziker et al, 2022; Hunziker & Stertz, 2022). All samples were analyzed by liquid chromatography tandem-mass spectrometry (LC-MS/MS) using a Thermo Scientific™ Orbitrap Fusion™ Tribrid™ mass spectrometer connected to an Easy-nLC 1000 HPLC system (Functional Genomics Center Zurich).

### Proteomic data analysis

Peptide identification, LFQ, and phosphorylation site determination were performed essentially as described (Hunziker & Stertz, 2022), but using the Progenesis QI for Proteomics software. Functional enrichment gene ontology terms (Aleksander *et al*, 2023; Ashburner *et al*, 2000) were determined with DAVID (Huang da *et al*., 2009b) using all human genes as background. Protein networks were created with STRING (Szklarczyk *et al*., 2015) and annotated in Cytoscape 3.9 (Shannon *et al*, 2003).

### Immunoprecipitations and western blot analyses

Treated cells were washed once with PBS and then lysed for 15 min on ice in lysis buffer (50mM Tris-HCl (pH7.4), 150mM NaCl, 1mM EDTA, 1% Triton X-100) supplemented with cOmplete Protease Inhibitor Cocktail (Roche) and PhosSTOP (Roche). Lysates were then cleared by centrifugation for 15 min at 14,000 rpm at 4°C and subjected to immunoprecipitation. Lysates were incubated with specific antibodies overnight at 4°C with rotation. 30 μl Protein G Dynabeads (Invitrogen) were then added and incubated for a further 30 min at room temperature with rotation. Beads were washed three times with lysis buffer. As required, samples were treated with λ- phosphatase (P0753S, NEB) according to the manufacturer’s instructions, and with recombinant Flag-IKKε (#31177, Active Motif) in kinase assay buffer (#9802, Cell Signaling) supplemented with 1 mM ATP for 30 min at 30 °C with shaking at 550 rpm. Immune complexes were eluted from beads by boiling for 10 min at 95°C in 1x Laemmli buffer (Bio-Rad), and were subjected to SDS-PAGE on Bolt™ Bis-Tris Plus Mini Protein Gels (4-12%), followed by western blotting. Primary antibodies used were: rabbit anti-STAT1-pY701 (#9167, Cell Signaling); mouse anti-STAT1 (sc-417, Santa Cruz); mouse anti-beta-actin (C- 4) (sc-47778, Santa Cruz); mouse anti-ANXA2-pY24 (sc-135753, Santa Cruz); mouse anti- ANXA2 (sc-28385, Santa Cruz); rabbit anti-STAT1-pS727 (#9177, Cell Signaling); rabbit anti-STAT1 (#14994, Cell Signaling); mouse anti-MX1 (Steiner & Pavlovic, 2020), mouse anti-RIG-I (AG-20B-0009-C100, Adipogen); rabbit anti-IRF9 (#D2T8M, Cell Signaling); rabbit anti-STAT2-pY690 (#88410, Cell Signaling); mouse anti-STAT2 (A-7) (sc-1668, Santa Cruz); rabbit anti-Flag (F7425, Sigma Aldrich); mouse anti-Flag M2 (F1804, Sigma Aldrich); rabbit anti-pSer 14-3-3 Binding Motif (14-3-3 PBM) (#9601, Cell Signaling); rabbit anti-PLEKHG3 (PA5-46053, Thermo Fisher Scientific); rabbit anti-HA (#3724, Cell Signaling); and mouse anti-HA (#2367, Cell Signaling). Secondary antibodies used were: IRDye 800CW goat anti-mouse IgG (LI-COR, #926–32210); IRDye 800CW goat anti-rabbit IgG (LI-COR, #926– 32211); IRDye 680CW goat anti-mouse IgG (LI-COR, #926–68070); and IRDye 680CW goat anti-rabbit IgG (LI-COR, #926–68071). Western blots were imaged on a LI-COR Odyssey XF Imager and western blot bands were quantified using the Image Studio Lite software.

### RT-qPCR

Total RNA was extracted using the ReliaPrep™ RNA Cell Miniprep System (Z6012, Promega) and 500 ng of RNA was reverse transcribed using SuperScript III Reverse Transcriptase (Thermo Fisher Scientific) according to the manufacturer’s instructions. Real-time PCR was performed using Fast EvaGreen qPCR Master Mix (Biotium) using the forward and reverse primers listed in **Dataset EV5** in the 7300 Real-Time PCR System (Applied Biosystems). Relative gene expression was calculated with the ΔΔCt method, using GAPDH for normalization.

### RNAi screening, reporter virus assays, and ISRE-GFP assays

The RNAi screens were conducted in an arrayed 96-well plate format, with each of the 45 factors of interest targeted independently by four different siRNAs (Qiagen; targets in **Dataset EV5**). To assess the impact of siRNAs on the antiviral activity of IFN-I, A549 cells were transfected with each siRNA for 48 h, followed by treatment with 10 ng/ml of IFNα2 for 16 h. IAV-*Renilla* and VSV-GFP reporter viruses were then used to assess the resulting antiviral effects. For IAV- *Renilla*, cells were infected at an MOI of 1 PFU/cell for 1 h at 37°C, then washed with PBS and overlaid with DMEM supplemented with 0.1% FBS, 100 U/ml penicillin, 100 μg/ml streptomycin, 0.3% BSA, 20 mM HEPES, and 1 μg/ml TPCK-trypsin (Sigma-Aldrich), along with 6 μM EnduRen™ Live Cell Substrate (Promega). Luminescence was then measured over 12 h using the EnVision Multilabel Reader (Perkin Elmer), and area under the curve (AUC) values were calculated from the relative light units (RLUs) using GraphPad Prism software. For VSV-GFP, cells were infected at an MOI of 1 PFU/cell for 1 h at 37°C, then washed with PBS and overlaid with Fluorobrite DMEM (Life Technologies) supplemented with 2% FBS, 100 U/ml penicillin, 100 μg/ml streptomycin, and 2 mM L-glutamine. GFP expression was monitored for up to 72 h using the IncuCyte ZOOM Live-Cell Imaging System (Sartorius), and Total Green Object Integrated Intensity (GCU x μm2/ well) values were used to calculate the AUC. To assess the impact of siRNAs on IFN-I signaling, A549/pr(ISRE).eGFP cells were transfected with each siRNA for 30 h, followed by treatment with 10 ng/ml of IFNα2. GFP expression was monitored for a further 72 h using the IncuCyte ZOOM, and AUC values were calculated as described above.

## Acknowledgements

We thank Rick Randall and Catherine Adamson (University of St Andrews, UK), as well as Peter Palese (Icahn School of Medicine, New York, USA), for kind gifts of cells and viruses. We also thank Feng-Qian Li & Ken-Ichi Takemaru, the Michael J Fox Foundation MJFF, Michael Yaffe, Jim Darnell, and Didier Trono for plasmids deposited at Addgene. Mass spectrometry experiments were performed at the Functional Genomics Center Zurich (FGCZ) of the University of Zurich and the ETH Zurich. The research leading to these results received funding from the Swiss National Science Foundation (grants 31003A_182464 and 310030_214957 to BGH) and the University of Zurich (grant FK-18-044 to IB). The funders had no role in study design, data collection, data interpretation, or the decision to submit the work for publication.

## Author contributions

Conceptualization: IB, ML and BGH; Methodology and Investigation: IB, ML, SF, SS and NT; Writing and Visualization: IB, ML, SF, SS and BGH; Supervision: BGH; Funding Acquisition and Project Administration: BGH.

## Supporting information

Expanded View Figures PDF Dataset EV1: Total Proteome Changes During Stimulation with IFNα, IFNβ, IFNω, IFNγ, and IFNλ. Dataset EV2: Phosphoproteome Changes During Stimulation with IFNα, IFNβ, IFNω, IFNγ, and IFNλ. Dataset EV3: Overlap of Proteomic and Phosphoproteomic Changes after Stimulation with different IFNs. Dataset EV4: RNAi Screening Control Data. Dataset EV5: Materials

## Data availability

The mass spectrometry proteomics data have been deposited to the ProteomeXchange Consortium via the PRIDE (Perez-Riverol *et al*, 2019) partner repository with the dataset identifier PXD029969. The data will be made available following peer-review.

## Expanded View Figure Legends

Figure EV1. **Temporal Changes to the Total Human Proteome and Phosphoproteome During IFN**γ **or IFN**λ **Stimulation**.

**A** A549 cells underwent overnight serum starvation, followed by stimulation with 10 ng/ml of IFNγ or IFNλ1 for the indicated durations. Subsequently, the cells were analyzed by western blotting with the specified antibodies. Data presented are representative of n=3 independent experiments.

**B-C** Numbers of distinct proteins (**B**) and phosphorylated proteins (**C**) significantly up- or down-regulated at the indicated time points following stimulation with IFNγ (yellow) or IFNλ1 (blue), relative to non-stimulated samples.

**D-G** Venn diagrams illustrating comparisons between significantly up- or down-regulated proteins (**D/E**) or phosphoproteins (**F/G**) in response to stimulation with IFN-I (IFNα2, IFNβ, and IFNω), IFN-II (IFNγ), or IFN-III (IFNλ1).

**H-J** Circos plots displaying phosphoproteins with significantly enhanced phosphorylation after 30 min (**H**), 90 min (**I**), or 240 min (**J**) following stimulation with different IFNs (IFNα2, IFNβ, IFNω, IFNγ, or IFNλ1) compared to non-stimulated samples. Black lines connect identical peptides present in the datasets from multiple IFNs.

Figure EV2. Unique Phosphoproteome Changes in Response to IFN**α**2, IFN**β**, IFN**ω**, or IFN**γ** Stimulation.

**A-D** Phosphoproteins uniquely regulated in response to IFNα2 (**A**), IFNβ (**B**), IFNω (**C**), or IFNγ (**D**), but not to any other IFN. No unique phosphoproteins were identified in response to IFNλ1 stimulation. Phosphoproteins were considered uniquely changed by one IFN when they exhibited a significant phosphorylation fold change > 2 after IFN stimulation for the specific IFN, but were not significantly changed with any of the other four IFNs tested, independent of fold change observed. Only log_2_(FC) > 1 and < -1 are depicted. IFN-I signaling interactors (INT) according to BioGRID and/or STRING are indicated with gray dots, and ISGs with a more than two-fold increase after IFN stimulation are indicated with blue dots. Names are written in bold where several distinct phosphorylated peptides were detected for that protein.

Figure EV3. Western Blot Validation of IFN-Stimulated Phosphorylations.

**A-C** A549 cells were subjected to overnight serum starvation, followed by stimulation with IFNα2 or IFNω (10 ng/ml) for the indicated durations. Subsequently, cell lysates were analyzed by western blot using the specified antibodies. Representative western blots as well as quantification of n=2 or n=3 independent experiments for STAT1-pS727 (**B**) and ANXA2-pY24 (**C**) are depicted. Bars represent mean values and standard deviations.

**D-E** Relative phosphopeptide intensities of the peptides containing the STAT1 S727 and ANXA2 Y24 phosphorylation sites in response to stimulation with the indicated IFN-Is at the indicated times post-stimulation. Data points represent means with standard deviations from the five independent replicates.

Figure EV4. Western Blot Quantifications from Experiments using Reconstituted STAT1-KO cells.

**A-B** Non-edited control A549 cells (ne) or STAT1-deficient (KO) A549 cells reconstituted with empty vector (EV), wild-type (WT) STAT1, or the indicated STAT1 mutants were serum-starved overnight and stimulated with IFNα2 (10 ng/ml) for 30 min. Total protein lysates were analyzed by western blotting with the indicated antibodies (see Figure 4B). Relative band intensities of western blot quantifications of independent replicates is depicted for (**A**) total STAT1 (relative to Actin) and (**B**) STAT1-pY701 relative to total STAT1. Bars represent mean values and standard deviations from n=3 independent experiments (each dot corresponds to one replicate). Significance was determined by two-way ANOVA and Tukey’s multiple comparisons test on log-transformed data (ns, non-significant).

**C-F** Cells described above were serum-starved overnight and stimulated with IFNα2 (10 ng/ml) for 6 h. Total protein lysates were analyzed by western blotting with specific antibodies (see Figure 4J) and band quantifications are shown for RIG-I (**C**), IRF9 (**D**) and STAT1 (**E**). Bars represent mean values and standard deviations from n=3 independent experiments (each dot corresponds to one replicate). Significance was determined by t-test on log-transformed data (*P ≤ 0.05; ***P ≤ 0.001; ns, non-significant).

Figure EV5. PLEKHG3 Knockdown Leads to Reduced ISG Expression but Does Not Affect Early STAT Tyrosine Phosphorylation.

**A-C** A549 cells were transfected for 30 h with individual siRNAs targeting *PLEKHG3*, *IRF9* as a positive control, or a non-targeting siRNA as a negative control, prior to stimulation with IFNα2 for 4 h (1 ng/ml). Total RNA was harvested, and expression of the indicated genes was assessed by RT-qPCR with specific primers. Bars represent mean values and standard deviations from n=3 independent experiments (each dot corresponds to one replicate). Significance was determined by two-way ANOVA with Dunnett’s multiple comparison test on log-transformed data (*P ≤ 0.05; ***P ≤ 0.001; ****P ≤ 0.0001; ns, non-significant).

**D** A549 cells were transfected for 30 h with individual siRNAs targeting *PLEKHG3*, *IFNAR2* as a positive control, or a non-targeting siRNA as a negative control, prior to stimulation with IFNα2 for 30 min (10 ng/ml). Total protein lysates were analyzed by western blot with the indicated antibodies. Data are representative of n=3 independent experiments.

Figure EV6. Knockdown of Individual 14-3-3s Differentially Impacts IFN Signaling and VSV-GFP Replication.

**A-B** A549 cells were transfected with a control siRNA, an siRNA targeting *IRF9*, or siRNA pools individually targeting 14-3-3 family members (14-3-3β, 14-3-3γ, 14-3-3ε, 14-3-3ζ, 14-3- 3η, 14-3-3θ, or 14-3-3σ). 48 h post transfection, cells were stimulated with IFNα2 (1 ng/ml) for 4 h. Some cells were harvested and *IFIT2* mRNA levels were determined by RT-qPCR

(**A**). Some cells were infected with VSV-GFP at an MOI of 1 PFU/cell, and GFP expression as a measure for viral replication was assessed at 24 hpi (**B**). Bars represent mean values and standard deviations from n=3 independent experiments (each dot corresponds to one replicate). Significance was determined by two-way ANOVA with Dunnett’s multiple comparison test on log-transformed data (*P ≤ 0.05; **P ≤ 0.01; ***P ≤ 0.001; ****P ≤ 0.0001; ns, non-significant).

