## Supplementary figures and images for "An Atlas of Protein Phosphorylation Dynamics During Interferon Signaling"

### Supplemental Figures

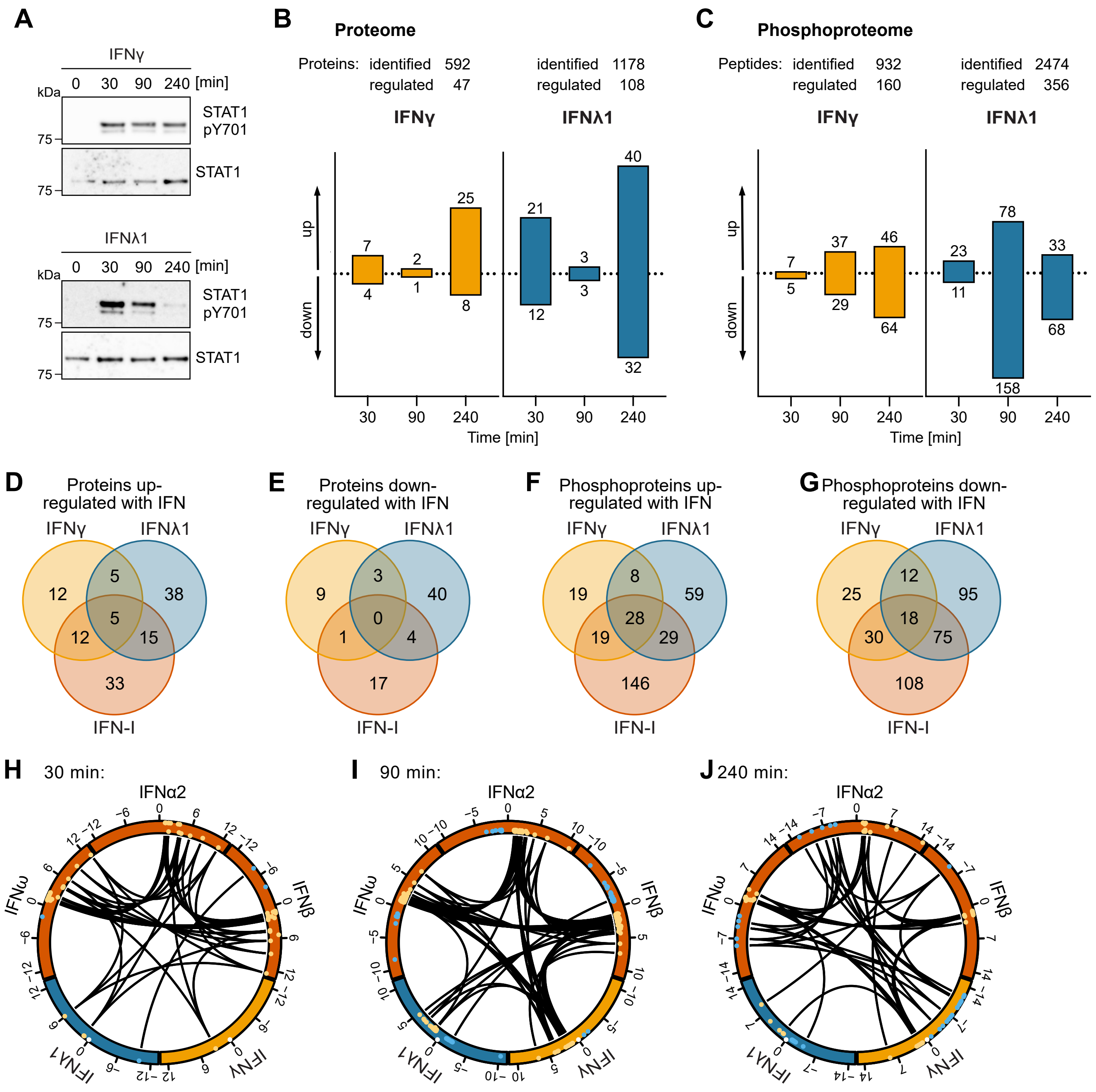

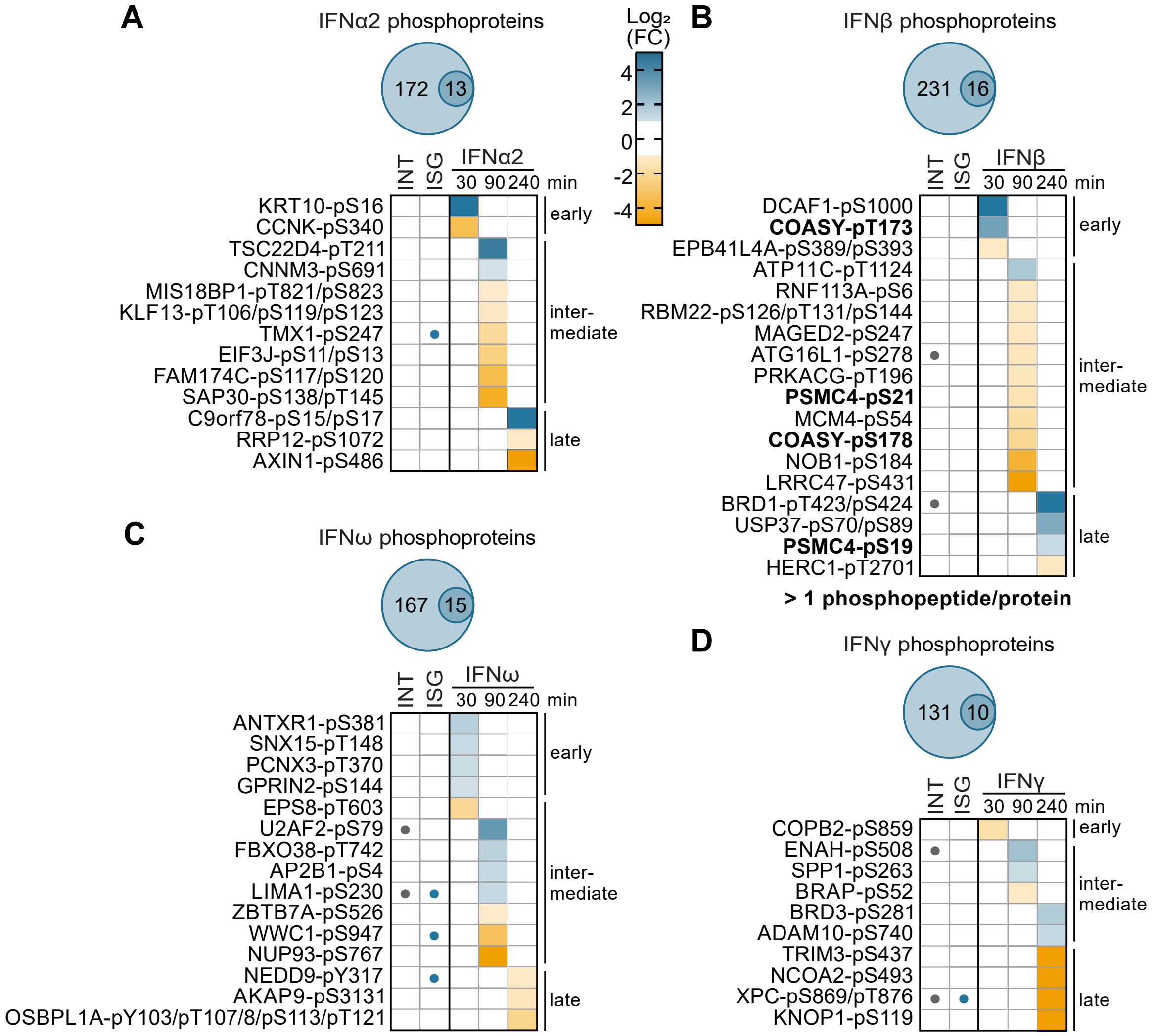

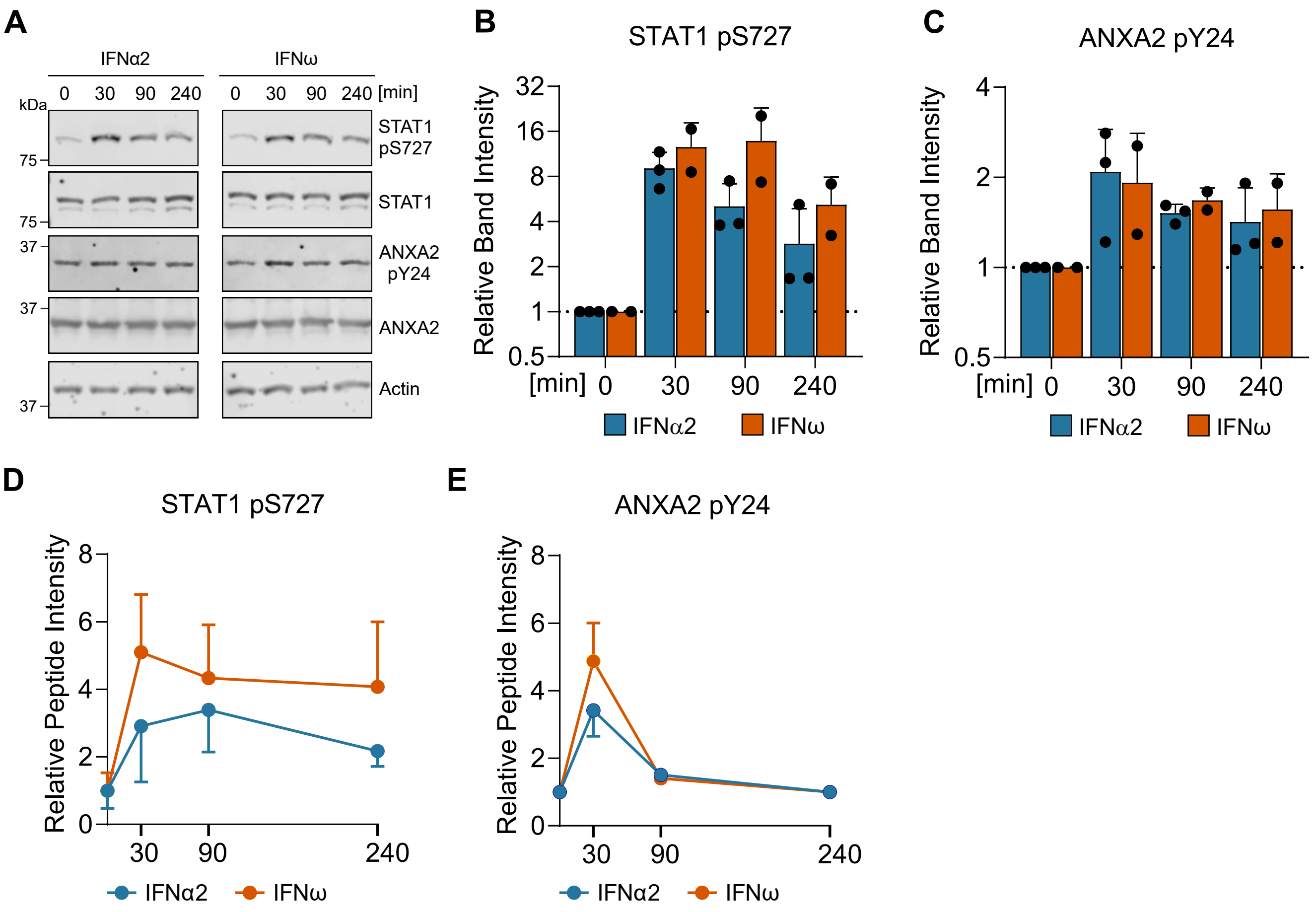

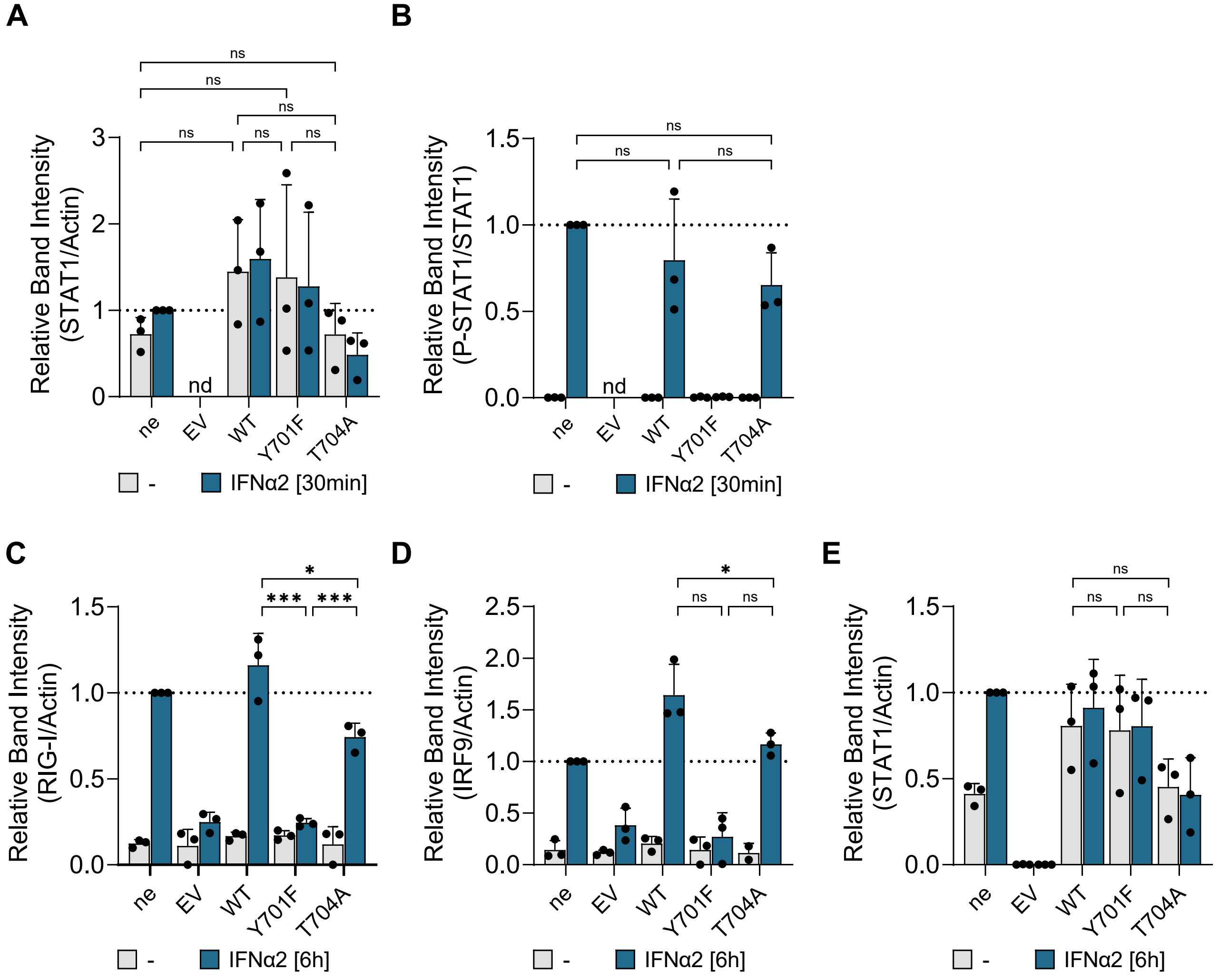

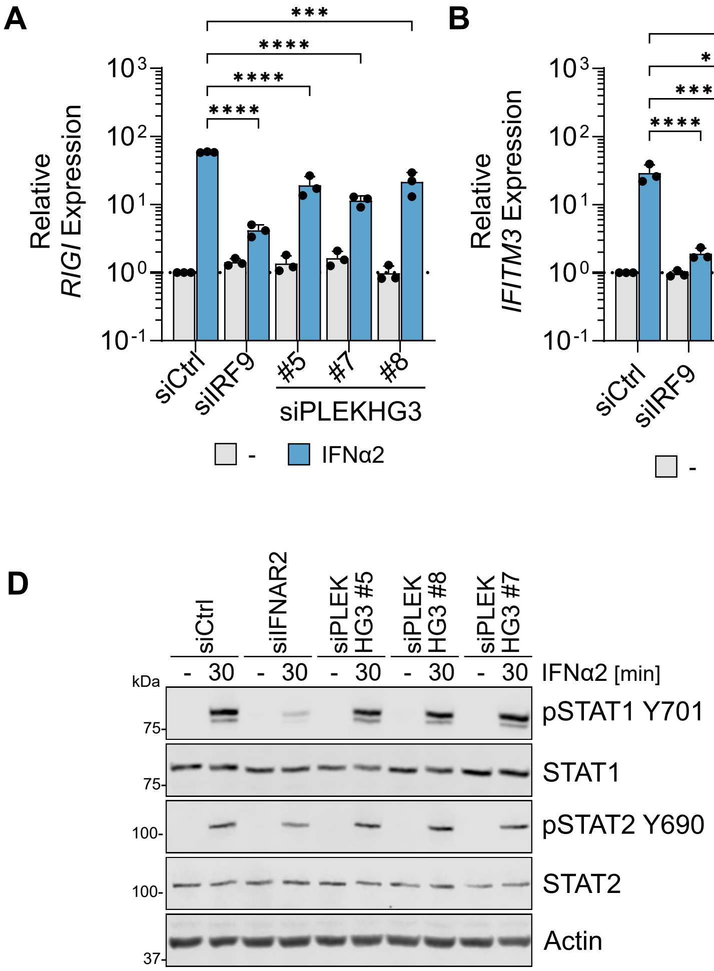

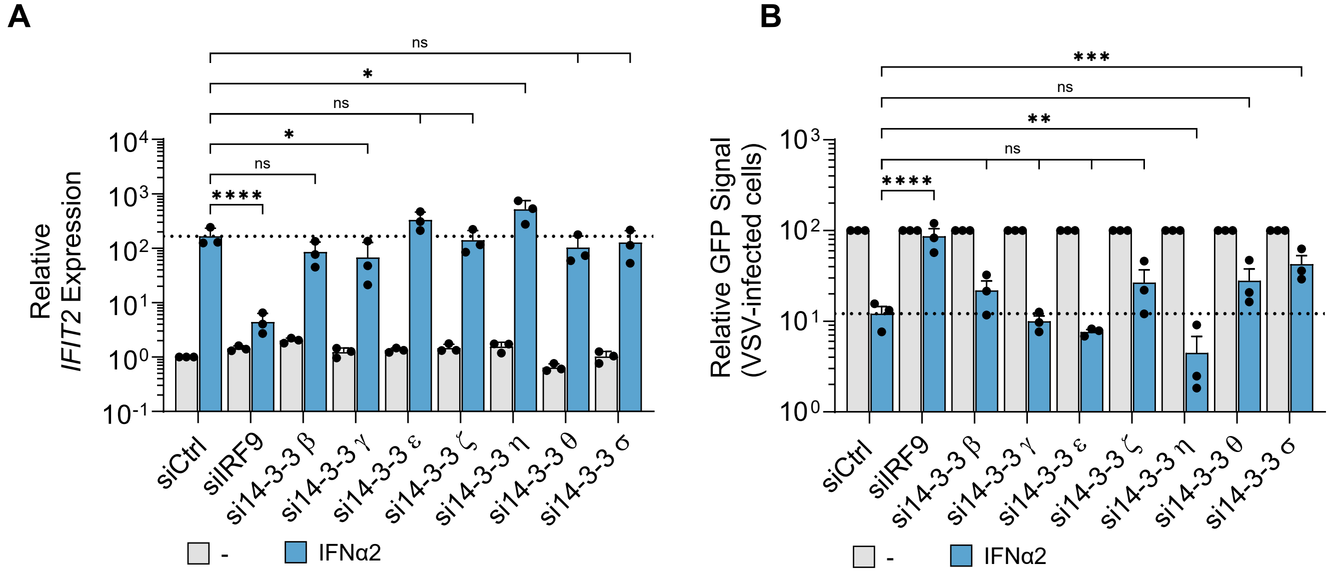
